# CD24-dependent innate immune checkpoint targeting chimera disrupts TGF-β-driven immunosuppression to enhance anti-tumor therapy

**DOI:** 10.64898/2026.09.24.754107

**Authors:** Ziyue Cui, Zhaoyi Lu, Mingzhu Pan, Yefan Hu, Mengchen Gao, Anzhe Ma, Yaqi Hong, Yuguo Zhang, Xi Chen, Bo Wang, Juan Zhang

**Author notes:** Corresponding author. (X.C.), (B.W.), (J.Z.). **One Sentence Summary:** A bifunctional CD24-driven innate immune checkpoint targeting chimera demonstrates superior tumor suppression via CD24-mediated TGF-β degradation.

## Abstract

The immunosuppressive tumor microenvironment (TME) poses formidable challenges for cancer immunotherapies. Among the complex TME landscape, the cytokine TGF-β is an attractive soluble target for reinvigoration of anti-tumor immunity due to its pivotal function in immune cell suppression, extracellular matrix remodeling, and epithelial-to-mesenchymal transition (EMT) promotion, while immune checkpoint receptors exemplified by PD-1 and CD24 are promising surface targets. Though adaptive immune checkpoint inhibition is well established in clinical practice, TGF-β blockade or innate immune checkpoint blockade is yet to be harnessed for tumor treatment. Here, leveraging CD24 overexpression on multiple tumor tissues and its unique recycling kinetics, we designed an innate Immune Checkpoint TArgeting Chimera (ICTAC) G01, a bifunctional CD24 antibody-TGF-β receptor fusion protein that disrupts TGF-β signaling by TGF-β degradation instead of simple blockade and simultaneously inhibits the “don’t eat me” CD24 signaling. We demonstrated that G01 effectively suppressed TGF-β-Smad signaling, reversed the EMT, and attenuated fibrotic collagen deposition. Additionally, it antagonized CD24 signaling to unleash robust antibody-dependent effector activity, further reprogramming the immunologically cold TME toward inflammatory phenotype. By selectively targeting malignant cells while sparing CD24-expressing B cells, G01 demonstrated a favorable safety profile. Notably, in patient-derived tumor fragments (PDTFs), G01 also inflamed the TME, underscoring its translational potential. Together, these results establish ICTAC as an effective strategy for tumor-specific receptor-mediated extracellular protein depletion, which supports its application for reversing TGF-β and innate immune checkpoint-mediated tumor immunosuppression by concurrent targeting of both surface and soluble tumor-promoting factors.

## INTRODUCTION

Cancer immunotherapies have revolutionized the immuno-oncology landscape by reinvigorating the anti-tumor capabilities of the immune system. Despite their success in certain tumor indications, the majority of patients with multiple types of solid tumors do not respond or develop resistance (*1–3*). The tumor microenvironment (TME) where different immunosuppressive factors including cytokines, immune checkpoints, and extracellular matrix barrier integrate to inhibit anti-tumor immunity is central to tumor immune escape (*4–6*). Among the complex cytokine network in the TME, the pleiotropic transforming growth factor-beta (TGF-β) is being scrutinized extensively thanks to its critical functions in immune regulation and tissue remodeling (*7*). TGF-β suppresses cytotoxic CD8^+^ and conventional CD4^+^ T cells by cooperating with transcription factor ATF1, promotes Tregs by inducing Foxp3 expression, and induces tolerogenic DCs by harnessing transcription factor ID1 (*8–10*). TGF-β also activates cancer-associated fibroblasts and promotes ECM remodeling, creating biophysical barriers for immune cell infiltration (*11–14*). Additionally, TGF-β has been demonstrated to promote epithelial-mesenchymal transition (EMT), which contributes to tumor metastasis. All these features make effective targeting of TGF-β a long-sought aim for cancer treatment.

With respect to immune checkpoints, multiple T cell-associated receptors such as PD-1, CTLA4, and LAG-3 have been carefully examined. Among these, PD-1 is the most well-established target for reinvigoration of anti-tumor immunity, with quite a few blocking antibodies approved in clinics. Interestingly, beyond these adaptive immune-related receptors, numerous innate immune checkpoints, e.g., CD47 and CD24 have been identified and are drawing much attention. These innate checkpoints function by interacting with specific receptors on myeloid lineage cells such as macrophages and dendritic cells to deliver “don’t eat me” signals. For instance, the two highly glycosylated checkpoints CD47 and CD24 signal via SIRPα and Siglec-10 on tumor-associated macrophages (TAMs), respectively, to inhibit their phagocytotic functions (*15–17*). Thus disrupting these interactions promises to promote tumor ingestion and antigen presentation for adaptive immune elicitation.

Clinically, co-upregulation of both TGF-β and immune checkpoints is often observed. Such co-elevation correlates with shortened patient survival and exacerbated T cell exhaustion across multiple high-incidence malignancies, underscoring the critical need for a dual-targeting strategy to augment tumor immunotherapy (*18*). Though promising, multiple anti-PD-1 antibody-TGF-β trap fusion proteins that concurrently target PD-1 and TGF-β have not shown clinical benefits (*19–24*). Dose-limiting cardiovascular toxicities and “sink effect” in which the vast reservoir of circulating and stroma-sequestered TGF-β rapidly saturates neutralizing agents have been identified to contribute to the low efficacy and severe side effects by these modalities (*25–30*). Interestingly, these fusion proteins all disrupt TGF-β signaling by blocking its interaction with TGF-β receptor, potentially indicating that TGF-β blockade only may not be sufficient. Importantly, to date, co-targeting of innate immune checkpoints and TGF-β remains to be explored.

Here, we developed a new immune checkpoint and immunosuppressive cytokine TGF-β co-targeting strategy named innate Immune Checkpoint TArgeting Chimera (ICTAC). Inspired by the unique rapid recycling of the innate immune checkpoint CD24 and multiple recently reported target protein degrading platforms, we designed G01, a bifunctional fusion protein composed of an anti-human CD24 monoclonal antibody fused with the extracellular domain (ECD) of TGF-β receptor 2 (TGFBR2). Unlike previously engineered TGF-β targeting proteins that block TGF-β signaling via neutralization, G01 performs such task via CD24-mediated TGF-β degradation, thus can effectively deplete TGF-β in the TME with no sink effect. Beyond TGF-β removal, G01 also disrupted CD24-Siglec-10 interaction to reinvigorate macrophage-dependent anti-tumor immunity. We showed that G01 disrupted TGF-β-Smad signaling to inhibit tumor proliferation and EMT, and to remodel the TME toward an immune inflammatory landscape. In PDTFs from

HNSCC patients, G01 also reactivated anti-tumor immune cells, suggesting its capability to remodel the human TME. Importantly, thanks to the fact that CD24 is expressed at low levels in normal cells except B cells but upregulated upon malignant transformation, and the unique binding profile of our CD24 antibody that specifically recognizes tumor-associated but not B cell-expressed CD24, G01 demonstrated a favorable safety profile, sparing B cells and erythrocytes. Therefore, our ICTAC molecule G01 offers a novel pathway for TGF-β blockade, highlighting its potential for precise manipulation of other extracellular proteins via a unique category of innate immune checkpoints. The concurrent targeting mode of both soluble and surface tumor evasive factors by ICTAC also enables tumor treatment by leveraging two complementary layers of tumor-specific clues.

## RESULTS

### Co-dysregulation of CD24 and TGF-β1 correlates with poor prognosis and T cell exhaustion

To assess whether concurrent targeting of CD24 and TGF-β1 is clinically supported, we first quantified *TGFB1* and *CD24* expressions in multiple human malignancies. Comprehensive analysis of the TCGA and GTEx datasets revealed that *CD24* and *TGFB1* were significantly upregulated in various solid tumors with high incidence, notably in LIHC and BRCA, compared to their respective adjacent normal tissues (Fig. 1A). Additionally, concurrent upregulation of these two markers (*CD24*^high^/*TGFB1*^high^) defined a patient subgroup with worse clinical outcomes. Specifically, this dual-high signature correlated with shortened overall survival (OS) in patients with BRCA (P = 0.0045) and LIHC (P = 0.079) (Fig. 1B). We further investigated how dysregulation of these two targets impacted immune infiltration in the TME. Gene set variation analysis (GSVA) revealed the synergistic effect of CD24 and TGF-β1 in shaping an immunosuppressive landscape. Compared to the *CD24*^low^/*TGFB1*^low^ subgroup, patients with *CD24*^high^/*TGFB1*^high^ signature exhibited significantly higher T cell exhaustion scores across both BRCA and LIHC cohorts (P < 0.0001) (Fig. 1C). Collectively, these findings confirm the CD24/TGF-β1 axis as a critical molecular node driving immune evasion and support their dual targeting as a promising route to reinvigorate anti-tumor immunity.

**Fig. 1.**
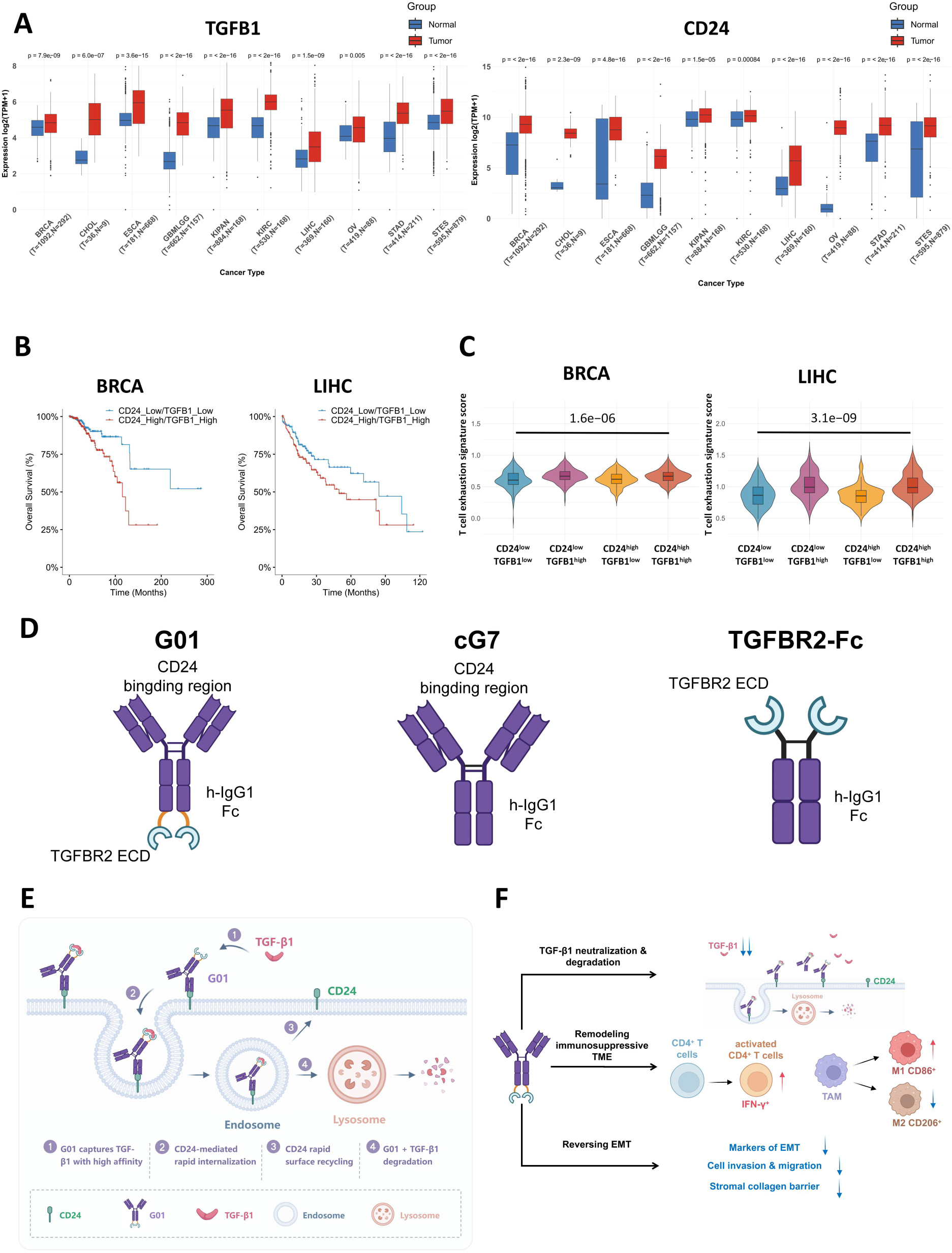
CD24 and TGF-β1 upregulation correlates with poor prognosis and immune exhaustion in multiple tumor indications and rationalizes their co-targeting via ICTAC. (A) *CD24* and *TGF-B1* expression in tumor and normal tissues from the TCGA database. Breast Invasive Carcinoma (BRCA), Cholangiocarcinoma (CHOL), Esophageal Carcinoma (ESCA), Glioblastoma and Lower Grade Glioma (GBMLGG), Pan-Kidney Cancer (KIPAN), Kidney Renal Clear Cell Carcinoma (KIRC), Liver Hepatocellular Carcinoma (LIHC), Ovarian Serous Cystadenocarcinoma (OV), Stomach Adenocarcinoma (STAD) and Stomach and Esophageal Carcinoma (STES). Statistical significance was assessed by unpaired, two-tailed Student’s t-test. **(B)** Overall survival of patients with BRCA (n = 245) and liver cancer (n = 120). Clinical metadata and log2-transformed transcriptomic data (log_2 [_TPM + 1]) for the TCGA-LIHC and TCGA-BRCA cohorts were retrieved from the UCSC Xena browser (https://xenabrowser.net). Patients are divided into *CD24* & *TGFB1* double high expression (red) and *CD24* & *TGFB1* double low expression (blue). Statistical analysis was performed with Log-rank test (P = 0.0045 and 0.079 for breast and liver cancer, respectively). **(C)** Calculation of T cell exhaustion scores using GSVA. RNA expression data were retrieved for the TCGA-LIHC and TCGA-BRCA cohorts, and the exhaustion scores were compared between BRCA (left) and LIHC (right) patients that were stratified by CD24 and TGFB1 expression. Statistical analysis was performed using paired t test. **(D)** Molecular design of CD24-TGF-β1 co-targeting ICTAC G01, anti-CD24 antibody cG7 and TGFBR2-Fc. **(E)** Mechanism of G01-driven lysosomal degradation of TGF-β1 via CD24-mediated internalization. **(F)** The multifaceted anti-tumor mechanisms of G01 examined in this study.

Importantly, CD24 has been shown to undergo constitutive endocytosis, laying the foundation for leveraging such feature for efficient degradation of extracellular target proteins by rapid internalization and lysosomal targeting. Inspired by multiple newly developed target protein degrading technologies, we designed a bifunctional ICTAC molecule G01 that not only exploits CD24 as a “shuttle” to deplete extracellular TGF-β1, but also blocks CD24 simultaneously. This ICTAC molecule is built by fusing a previously identified high-affinity anti-CD24 monoclonal antibody cG7 with the ECD of human TGF-β receptor II (TGFBR2) (Fig. 1D). We anticipate that this fusion protein, capable of simultaneously capturing cell-surface CD24 and soluble TGF-β1, would facilitate the rapid internalization and subsequent lysosomal degradation of TGF-β1 within the TME (Fig. 1E). Additionally, given the critical functions of CD24 and TGF-β1 in multiple immunosuppressive signaling pathways, we hypothesize that G01 will exert potent anti-tumor efficacy through multifaceted mechanisms, ranging from direct innate immune checkpoint blockade to the dismantling of the stromal collagen barrier (Fig. 1F). To investigate the potential synergy of dual CD24 and TGF-β1 targeting, the parental CD24 antibody cG7 and the TGFBR2-Fc fusion protein were also included in all studies hereafter.

### G01 effectively recognizes CD24 and TGF-β1 and spares B cells

We first produced G01 through transient expression in HEK293F cells and performed comprehensive biophysical and biochemical characterizations. Size-exclusion high-performance liquid chromatography (SEC-HPLC) revealed that G01 exhibited high monodispersity and purity (>95%) with negligible aggregation or degradation (Fig. S1A). Its structural integrity was further validated by SDS-PAGE, which displayed single migration bands under non-reducing conditions. Under reducing conditions, the expected heavy and light chains were observed for G01, confirming the correct assembly of the bifunctional molecule (Fig. S1B).

Biolayer interferometry (BLI) was used to determine the binding kinetics of G01 to human CD24 and TGF-β1. G01 preserved high-affinity binding of its parent cG7 to hCD24 (K_D o_f 9.08 and 8.18 nM, respectively; Fig. S1C). As expected, the affinity of G01 to hTGF-β1 was in the same sub-nanomolar range as that of TGFBR2-Fc (K_D o_f 0.776 and 0.197 nM, respectively). Since pan-TGF-β blockade is notoriously associated with dose-limiting cardiotoxicity due to the perturbation of hTGF-β2 and hTGF-β3 that are essential for cardiac homeostasis, we assessed the recognition of these two isoforms by G01. Notably, G01 exhibited an approximately 3-fold lower affinity for hTGF-β2 compared to hTGF-β1, characterized by rapid dissociation with the hTGF-β2 isoform (Fig. S1D). Though displaying comparable affinity for hTGF-β3 as for hTGF-β1, the physiological abundance of hTGF-β3 is substantially lower in vivo. Therefore, this isoform-biased binding profile may mitigate the cardiotoxicities resulted from non-selective TGF-β inhibition.

At the cellular level, G01 recognized CHO cells overexpressing human CD24 (CHO-hCD24) but not WT CHO or CHO overexpressing murine CD24 (CHO-mCD24) (Fig. S1E). Furthermore, G01 effectively associates with endogenous CD24 on human hepatocarcinoma cell lines (PLC/PRF/5 and BEL-7402) (Fig. S1F), and its capability of concurrent CD24 and TGF-β1 binding validated the functional integrity of the dual-targeting design (Fig. S1G). Importantly, a non-negligible safety concern of CD24 targeting is that CD24 is physiologically expressed on normal plasma cells. To evaluate potential on-target, off-tumor toxicities, we assessed G01 binding to primary plasma cells isolated from healthy human peripheral blood mononuclear cells (PBMCs). Interestingly, unlike the control antibody SN3 that robustly recognizes plasma cells, G01 exhibited very low binding to plasma cells, indicating its superior tumor selectivity, which resulted from its parent cG7 that also preferentially targeted tumor cells instead of B cells (Fig. S1H). This tumor-selective binding profile of G01 supports its favorable safety and tolerability.

Finally, to ensure that C-terminal fusion of the TGFBR2-ECD did not sterically hinder the IgG1 Fc domain, we measured G01 binding to the high-affinity variant of the activating Fc receptor CD16a (V158). Both G01 and cG7 successfully engaged CD16a-overexpressing Jurkat cells with comparable apparent affinities (EC_50 v_alues of 38.4 and 50.9 nM by flow cytometry, respectively; Fig. S1I), suggesting the retention of Fc effector function.

Together, these data confirm that G01 is a structurally stable, bifunctional molecule capable of simultaneously targeting hCD24 and hTGF-β1.

### G01 hijacks CD24 endocytosis to degrade extracellular TGF-β1

While all current antibody-TGF-β trap fusion proteins disrupt TGF-β signaling by blocking its interaction with TGF-β receptor, our ICTAC molecule G01 is expected to directly degrade TGF-β1 in TME by harnessing the internalization machinery of CD24. To determine whether G01 can deplete extracellular TGF-β1, we first examined its internalization kinetics by incubating G01 with CD24 positive tumor cells under two different conditions. Upon transient G01 exposure where excess unbound molecules were removed from the culture, flow cytometric analysis revealed its rapid internalization at 37°C, which achieved 84.9% internalization within 30 minutes and plateaued at >90% by 90 minutes (Fig. 2, A-B). This is consistent with the internalization kinetics of its parent cG7 (Fig. S2, A-B). As a reference, antibodies targeting other well-characterized tumor-specific receptors, such as Trop2 or HER2, showed much slower internalization (Fig. S2, C-F). Interestingly, when excess G01 was not washed away during the incubation which mimics sufficient supply of drug molecules after dosing in vivo, the fluorescence intensity of surface-bound G01 remained almost stable, with only slight decrease over time, suggesting that surface CD24 is not depleted by antibody-mediated internalization (Fig. 2C). Additionally, live-cell imaging using a pH-sensitive fluorescent probe (pHrodo) confirmed this rapid endocytic process. After a 2-hour incubation, both cG7 and G01 robustly accumulated within the acidic endolysosomal compartments, whereas TGFBR2-Fc failed to internalize, confirming that internalization is indeed CD24-dependent (Fig. 2D).

**Fig. 2.**
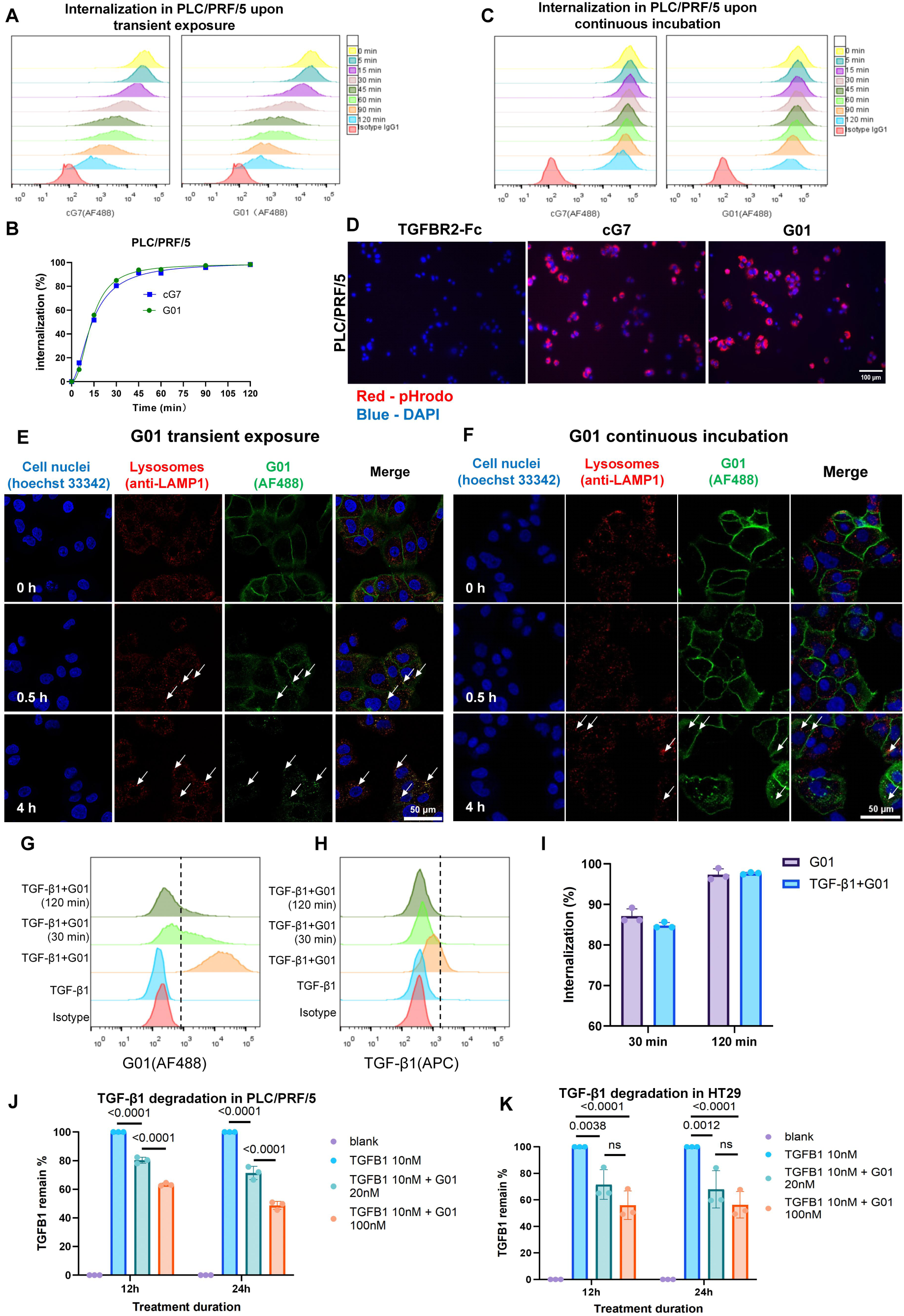
G01 drives rapid internalization and continuous lysosomal degradation of TGF-β1. **(A)** Surface G01 and cG7 fluorescence change on PLC/PRF/5 under transient exposure that remove excess unbound molecules. **(B)** Time-dependent internalization of G01 and cG7 in PLC/PRF/5 under transient exposure. Internalization (%) was calculated based on the mean fluorescence intensity (MFI) using the formula: [MFI (0 h) − MFI (n h)] / [MFI (0 h) − MFI (isotype)] × 100%. **(C)** Surface G01 and cG7 fluorescence change on PLC/PRF/5 under continuous incubation with the respective molecule without washing. **(D)** Live-cell imaging of G01 lysosomal entry. Representative fluorescence images of PLC/PRF/5 cells following a 2-hour incubation with pHrodo-conjugated cG7, G01, or TGFBR2-Fc. Scale bar, 100 μm. **(E and F)** Confocal imaging of the spatiotemporal trafficking of G01 to lysosomes in PLC/PRF/5 cells under (E) transient exposure or (F) continuous incubation of G01. Scale bar, 50 μm. **(G to I)** G01-mediated TGF-β1 internalization in engineered MC38-hCD24 cells detected via flow cytometry. Representative histograms of G01 (G) and TGF-β1 (H) levels after different incubation times were shown. (I) Internalization (%) was calculated based on the MFI using the formula: [MFI (0 h) - MFI (n h)] / [MFI (0 h) - MFI (isotype)] × 100%. (n = 3, data represent mean ± SEM). Statistical significance was determined by one-way ANOVA with Tukey’s post hoc test. **(J and K)** G01-mediated depletion of extracellular TGF-β1 in PLC/PRF/5 (J) and HT29 cells (K) quantified by ELISA. Data are representative of three independent experiments, and presented as mean ± SEM. Statistical significance was determined using two-way analysis of variance (ANOVA).

We mapped the intracellular trafficking of G01 using confocal microscopy. Under transient exposure, G01 was initially restricted to the plasma membrane. By 0.5 hour, its intracellular fluorescence intensified concurrently with a reduction in membrane signals; by 4 hours, G01 surface signals almost disappeared, with the intracellular G01 molecules translocated to and colocalized with LAMP1-positive lysosomes (Fig. 2E). Strikingly, under continuous incubation where excess G01 was not removed, its surface fluorescence remained robust, while the intracellular lysosomal accumulation steadily increased, again proving surface CD24 is not exhausted (Fig. 2F). Together with the flow cytometric analysis, these results indicate that CD24 functions as an efficient shuttle to route extracellular antibodies into the lysosomes.

To determine whether G01 can effectively drag its payload into tumor cells, we evaluated the internalization of the pre-formed G01/TGF-β1 complexes in CD24-positive cells. Upon incubation, these cells exhibited robust double-positive signals for G01 (AF488) and His-tagged TGF-β1 (APC). After incubation at 37°C, G01 and TGF-β1 fluorescence decayed concurrently, dropping to background levels by 120 minutes (Fig. 2, G-H). The internalization rate of the complex reached ∼85% at 30 minutes and ∼98% at 120 minutes, indicating that G01 can indeed drive effective TGF-β1 internalization (Fig. 2I).

We quantified TGF-β1 degradation by G01 in a co-incubation assay where CD24-expressing tumor cells, exogenous TGF-β1 and G01 were added together. ELISA measurements of the culture supernatants revealed that G01 rapidly depleted TGF-β1 in both PLC/PRF/5 and HT29 cultures in a time- and dose-dependent manner, achieving at least 50% clearance by 24 hours (Fig. 2, J-K). Together, these results establish that G01 acts as a targeted molecular scavenger, capturing and delivering extracellular TGF-β1 to lysosomes for degradation via CD24-mediated endocytosis, thereby promising to deplete TGF-β1 in the TME.

### G01 reverses immunosuppression and inhibits EMT by blocking TGF-β signaling

Having established that G01 triggers TGF-β1 degradation via CD24-mediated internalization, we next evaluated whether this novel mode of TGF-β1 blockade translates into superior inhibition of TGF-β1-driven immunosuppressive pathways compared to conventional neutralization strategies. We first assessed how G01 impacts canonical TGF-β1 signaling. In a reporter assay, G01 robustly suppressed TGF-β/Smad signaling with an IC_50 o_f 0.532 nM, whereas cG7 alone showed minimal effect (Fig. 3A). Notably, G01 exhibited more profound blockade of TGF-β signaling than TGFBR2-Fc, indicating that the fused TGFβRⅡ domain sequesters TGF-β1 more efficiently when coupled with the targeting antibody.

**Fig. 3.**
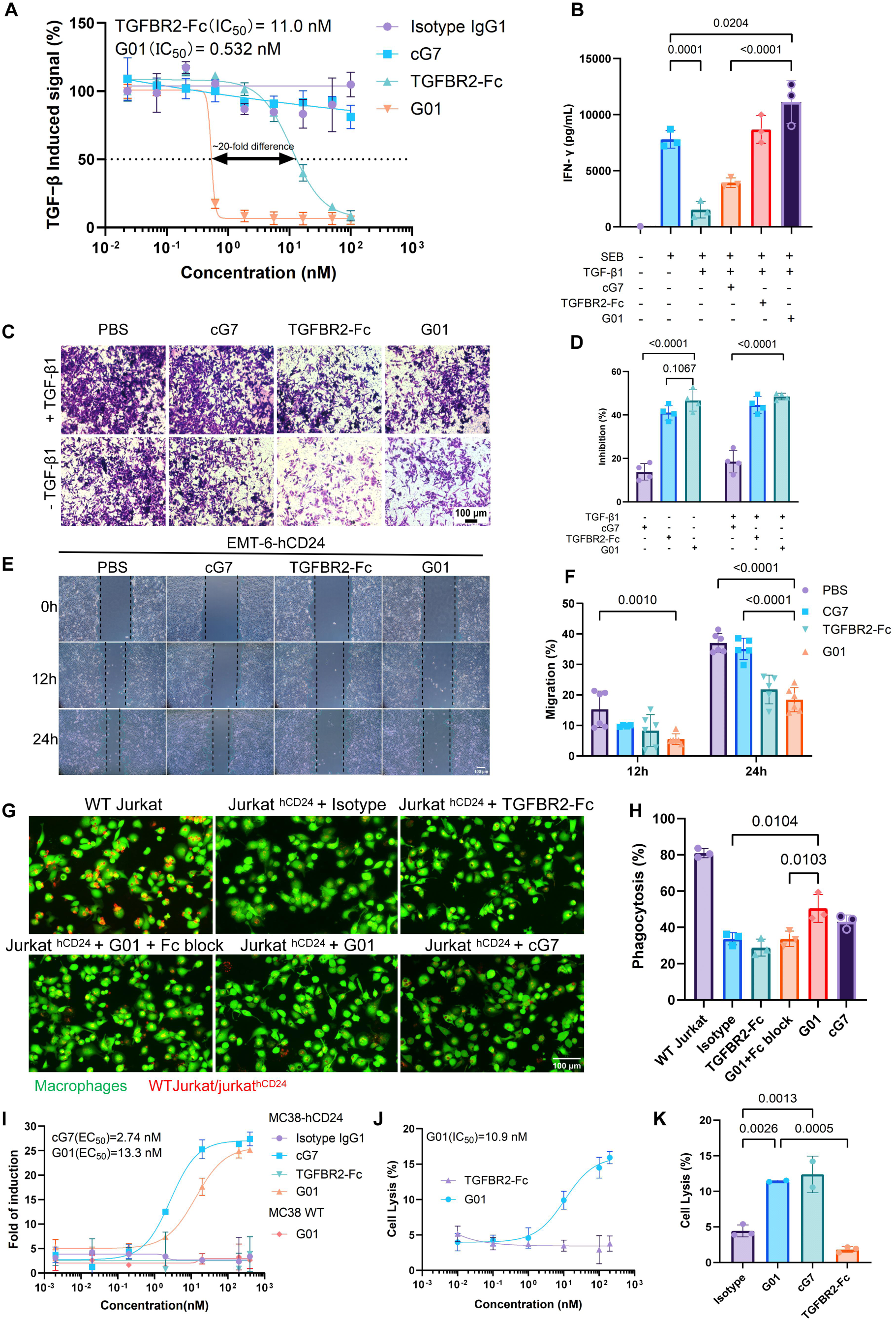
G01 disrupts TGF-β1-mediated suppressive signaling and triggers robust Fc-dependent cellular cytotoxicity and phagocytosis. **(A)** Dose-response plots showing the effect of G01, cG7, or TGFBR2-Fc on TGF-β/Smad signaling in HEK293/SBE-luc luciferase reporter cells (n = 3, data represent mean ± SD). **(B)** IFN-γ levels in the supernatant measured by ELISA after stimulation of human PBMCs with Staphylococcal enterotoxin B (SEB) and TGF-β1 together with the indicated drugs for 4 days (n = 3, data are representative from three independent experiments, and presented as mean ± SD). Statistical significance was determined by one-way ANOVA with multiple comparisons. **(C and D)** Representative images (C) and inhibition rate analysis (D) of Huh7 cell invasion (n = 4, data represent mean ± SD). Statistical significance was determined by one-way ANOVA with Tukey’s post hoc test. Scale bar, 100 μm. **(E and F)** Representative images (E) and quantification of the migration rate (F) from the scratch wound healing assay using EMT6-hCD24 cells (n = 5, data represent mean ± SD). Scale bar, 100 μm. Statistical significance was determined using two-way ANOVA. **(G and H)** G01 overrides CD24-mediated “don’t eat me” signal to promote ADCP. THP-1-derived macrophages (labeled with Calcein AM, green) were co-incubated with WT Jurkat or Jurkat-hCD24 cells (labeled with TraKine™, red) under various treatments for 6 hours. Shown are representative fluorescence images (G) and the calculated phagocytosis fraction (H), defined as the percentage of macrophages that engulfed target cells (n = 3, data represent mean ± SD). Statistical significance was determined by one-way ANOVA with Tukey’s post hoc test. Scale bar, 100 μm. **(I)** Quantification of ADCC via the reporter assay. MC38 or MC38-hCD24 target cells were pre-incubated with isotype IgG1, G01, cG7, or TGFBR2-Fc for 1 hour, followed by a 6-hour co-culture with Jurkat-Lucia™ NFAT-CD16 effector cells at an effector-to-target (E:T) ratio of 10:1. ADCC was evaluated by luciferase activity in the supernatant (n = 2, data represent mean ± SD). **(J and K)** PLC/PRF/5 tumor cell lysis by PBMCs upon different treatments for 4 hours at an E:T ratio of 20:1, evaluated by LDH release. (J) Dose-response curves of PLC/PRF/5 cytotoxicity comparing G01 and TGFBR2-Fc (n = 3, data represent mean ± SD). (K) Cytotoxicity comparison of isotype, cG7, TGFBR2-Fc, and G01 at a saturating concentration of 100 nM ((For isotype and TGFBR2-Fc group, n = 3; for cG7 and G01 group, n = 2; data represent mean ± SD). Statistical significance was determined by one-way ANOVA with multiple comparisons.

We evaluated whether G01 can reverse TGF-β1-mediated immunosuppression. Human PBMCs were stimulated with SEB in the presence of different treatments. TGF-β1 substantially blunted SEB-induced IFN-γ release (Fig. 3B). While TGFBR2-Fc restored IFN-γ secretion to baseline SEB levels and cG7 offered minimal activation, G01 triggered robust IFN-γ release that significantly exceeded the levels induced by SEB alone (p=0.0204). This synergistic effect indicates that G01 reverses TGF-β1-mediated immunosuppression via its trap domain and concurrent CD24 targeting that amplifies effector immune responses.

We investigated the impact of G01 on tumor phenotypic plasticity and motility. TGF-β1 exposure induced profound EMT morphological changes in PLC/PRF/5 cells, characterized by the loss of tight junctions, assumption of a loose mesenchymal phenotype, and pseudopodia formation (Fig. S3A). Both G01 and TGFBR2-Fc, but not cG7, effectively reversed these transitions. TGF-β1-potentiated cellular migration and invasion were also markedly abrogated by G01 in scratch wound and trans-well assays (Fig. 3, C-F, and Fig. S3, B-C). Importantly, while G01 and TGFBR2-Fc exhibited comparable efficacy in neutralizing high-dose exogenous TGF-β1, G01 demonstrated superior inhibition of BEL-7402 tumor cell invasion under basal conditions (without exogenous TGF-β1 at 12 hours) (Fig. S3C).

These results suggest that G01 can promote multiple anti-tumor mechanisms upon TGF-β1 degradation.

### G01 possesses CD24-dependent ADCP and ADCC effector functions

Beyond modulating the immunosuppressive TME via TGF-β1 neutralization, the intact IgG1 Fc domain should impart G01 Fc-mediated effector functions, including ADCP and ADCC. To evaluate its ADCP activity, we used THP-1-derived macrophages as effectors, which were polarized to M2-like phenotype to mimic TAM in the TME. Successful M2 polarization was validated by the robust upregulation of surface CD206 and Siglec-10 (Fig. S4A). Using Jurkat cells engineered to overexpress human CD24 (Jurkat-hCD24) as target cells with E:T of 1:2, we observed that CD24 expression markedly suppressed phagocytosis by Siglec-10^+^ macrophages (from 80.9% to 33.2%, WT Jurkat v.s. isotype) (Fig. 3G-H). Treatment with either G01 or cG7 partially restored phagocytosis, while TGFBR2-Fc which lacks CD24-targeting capability failed to enhance phagocytosis, demonstrating that G01-mediated ADCP requires CD24-dependent effector-target bridging. Furthermore, G01 phagocytosis was substantially abrogated upon treatment with a commercial Fc receptor blocking reagent, confirming its dependence on FcγR engagement. This potent phagocytic activity was further confirmed using human PBMC-derived Siglec-10^+^ macrophages, where G01 promoted tumor cell engulfment (17.2%) compared to that of an isotype control (4.03%), closely mirroring the efficacy of cG7 (23.6%) (Fig. S4B-C). Collectively, G01 relieves CD24-Siglec-10-mediated “don’t eat me” signaling and triggers Fc-dependent phagocytosis.

We next examined the ADCC activity of G01. Both G01 and cG7 efficiently bind human CD16a expressed on engineered Jurkat cells (Fig. S1I). We then employed a Jurkat-Lucia NFAT-CD16 reporter cell line to confirm FcγRIIIa activation upon G01 treatment. When co-cultured with MC38-hCD24 target cells at an effector-to-target ratio of 10:1, both G01 and cG7 engaged CD16a and induced comparable levels of luciferase reporter activation. As expected, G01 showed no activity against wild-type MC38 cells, underscoring the requirement of CD24 engagement for ADCC (Fig. 3I). Similarly, in an LDH release assay using human PBMCs as effector cells (E:T of 10:1), G01 efficiently lysed target tumor cells with 10.9 nM EC_50 (_Fig. 3J), and achieved comparable lysis to that of cG7 at 100nM drug concentration (Fig. 3K). Interestingly, independent of the Fc effector functions, G01 also exhibited a direct, CD24-dependent inhibition of tumor cell proliferation (Fig. S4D).

Together, these data indicate that G01 elicits potent effector functions against CD24-expressing tumor cells and suppress tumor cell growth by blocking CD24.

### G01 demonstrates potent anti-tumor efficacy with favorable safety profiles in vivo

We evaluated the in vivo efficacy of G01 in multiple mouse tumor models. Given that CD24 and TGF-β1 are co-dysregulated in liver cancer (Fig. 1A-C), we first utilized the human LIHC cell line BEL-7402, characterized by moderate-to-high CD24 expression and robust tumorigenicity, to establish a nude mouse xenograft model. G01 treatment elicited robust tumor regression, outperforming both the cG7 antibody and the standard-of-care sorafenib treatments (Fig. 4A and Fig. S5, A-B). Specifically, G01 achieved 62.4% tumor growth inhibition, which was superior to cG7 (25.6%) and slightly exceeded that of sorafenib (53.9%) (Fig. 4B). This robust anti-tumor efficacy translated into significant survival benefit; while all mice in the cG7 and sorafenib groups succumbed to tumor progression within 59 days, all G01-treated mice survived beyond 56 days (Fig. 4C). IHC analysis of Ki-67 confirmed that the anti-proliferative activity of G01 was comparable to that of sorafenib (26.1% vs. 31.8% Ki-67^+^ cells in the tumor sections, respectively) (Fig. 4, D-E). Notably, G01 exhibited good safety profiles. Throughout the treatment, G01-treated mice maintained steady body weight gains (Fig. 4F). Moreover, kidney weights of G01-treated mice remained indistinguishable from those of healthy controls, highlighting its excellent in vivo tolerability (Fig. 4, G-H).

**Fig. 4.**
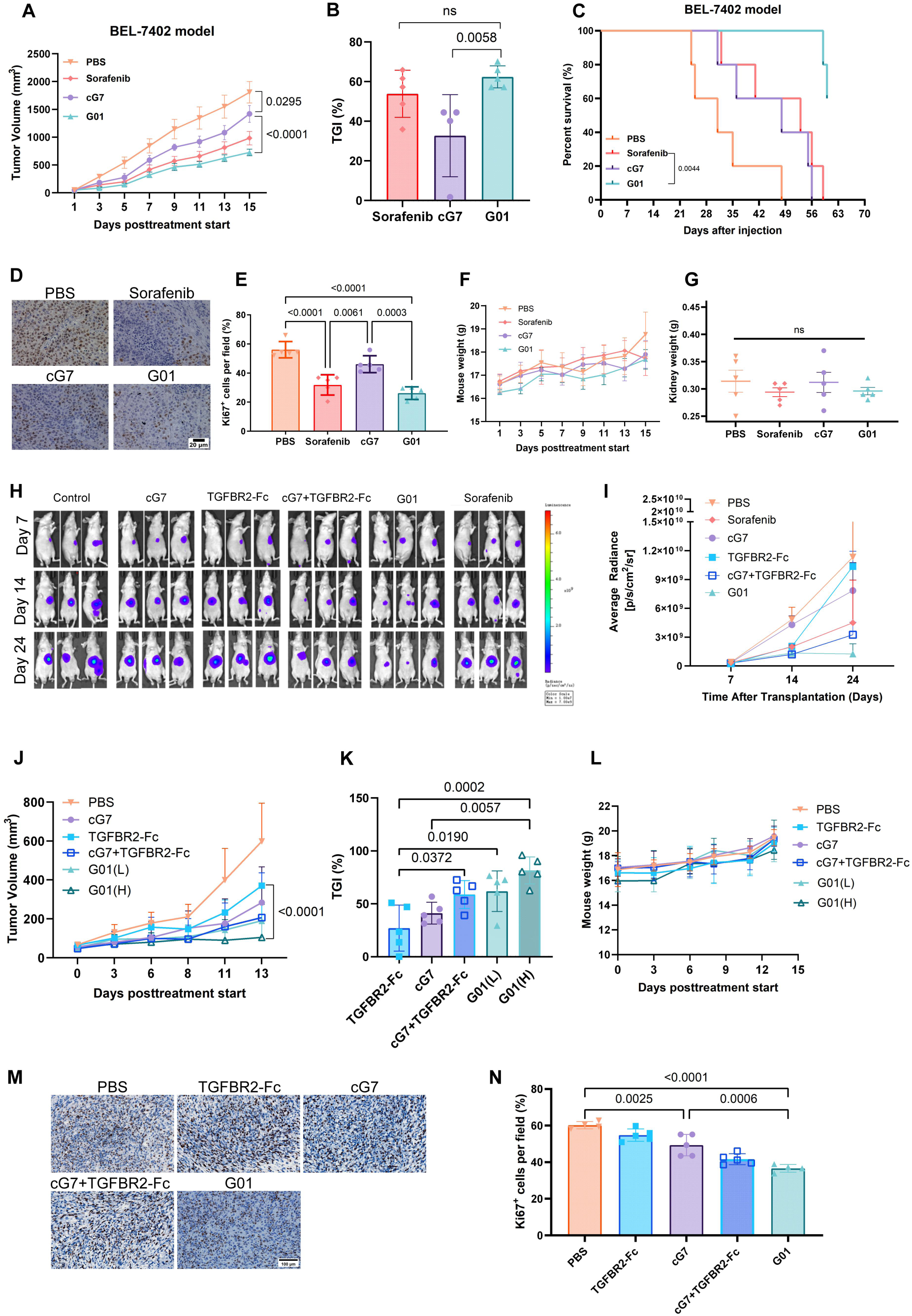
G01 demonstrates robust antitumor efficacy and favorable safety profile across various preclinical models. **(A to G)** BALB/c-Nude mice bearing established BEL-7402 human LIHC xenograft tumors were randomized (n = 5 per group) and treated intraperitoneally every two or three days for a total of six doses with G01 (10 mg/kg) or equimolar doses of cG7 (8.3 mg/kg). Sorafenib (20 mg/kg, daily oral gavage) served as a positive control, and PBS was used as a vehicle control. Treatment began when tumor volumes reached 50-100 mm^3^, and tumor volumes and body weights were monitored every two or three days. **(A to B)** Tumor growth kinetics (A) and tumor growth inhibition calculated with the corresponding endpoint tumor weights (B). Data are presented as mean ± SD, statistical significance was determined by two-way ANOVA (A) or one-way ANOVA with Tukey’s post hoc test (B). **(C)** Kaplan-Meier survival analysis of BEL-7402 tumor-bearing mice. Statistical significance was determined by Log-rank (Mantel-Cox) test. **(D and E)** Representative immunohistochemical (IHC) images (D) and quantification (E) of the proliferation marker Ki-67 in excised BEL-7402 tumors. Scale bars, 20 μm. Data represent mean ± SD. Statistical significance was determined by one-way ANOVA with Tukey’s post hoc test. **(F)** Body weights of BEL-7402 tumor-bearing mice across treatment groups. Data represent mean ± SEM. **(G)** Safety evaluation. Kidney weights demonstrate that G01 and cG7 are well-tolerated with negligible renal toxicity. Data represent mean ± SD. Statistical significance was determined by one-way ANOVA with Tukey’s post hoc test. **(H and I)** Antitumor efficacy in an orthotopic liver tumor model. PLC/PRF/5-luc cells were surgically implanted into the left liver lobe of nude mice. Treatments were initiated on day 7 post-implantation, after confirmation of tumor engraftment. Mice bearing established tumors were randomized (n = 3 per group) and treated intraperitoneally every two or three days for a total of six doses with G01 (10 mg/kg) or equimolar doses of cG7 (8.3 mg/kg), TGFBR2-Fc (4.6 mg/kg), or their combination. Tumor growth was monitored using bioluminescence imaging, and representative images (H) and quantification of luminescent signals (I) are shown for days 7, 14, and 24. Data represent mean ± SD, statistical significance was determined by two-way ANOVA. **(J to N)** C57BL/6J mice bearing MC38-hCD24 colorectal tumors were randomized and treated intraperitoneally every two or three days for a total of six doses with G01 at high (10 mg/kg) or low (3 mg/kg) doses, or equimolar doses of cG7 (8.3 mg/kg), TGFBR2-Fc (4.6 mg/kg), or their combination. Tumor volumes and body weights were monitored every two or three days. n = 8 for PBS, cG7, cG7 + TGFBR2-Fc and G01 (H) groups; n = 7 for TGFBR2-Fc group; n = 5 for G01 (L) group. Tumor growth kinetics (J) and endpoint tumor inhibition (K). Data represent mean ± SD, and statistical significance was determined by two-way ANOVA (J) or one-way ANOVA with multiple comparisons (K). **(L)** Body weights of MC38-hCD24 tumor-bearing mice throughout the treatment. Data represent mean ± SD. **(M and N)** Representative IHC staining and quantification of Ki-67 expression in MC38-hCD24 tumor tissues to evaluate cellular proliferation. Data represent mean ± SD, and statistical significance was determined by one-way ANOVA with Tukey’s post hoc test.

We also employed an orthotopic liver tumor model that recapitulates human liver vascularity and the physiological microenvironment to evaluate the anti-tumor efficacy of G01. PLC/PRF/5 cells that exhibit high CD24 expression and TGF-β sensitivity were engineered to express firefly luciferase (PLC/PRF/5-Luc) for in vivo tracking of tumor growth, which proliferated at the same rate as the parental cells in vitro (Fig. S5C). Longitudinal bioluminescence imaging revealed that while TGFBR2-Fc and cG7 provided marginal-to-modest delays in tumor progression, G01 resulted in remarkable tumor growth suppression, even superior to sorafenib (Fig. 4, I-J).

We next used an immunocompetent syngeneic model to investigate the anti-tumor efficacy of G01. Given that G01 does not cross-react with murine CD24, we generated a stable MC38-hCD24 cell line. Though displaying similar proliferation kinetics in vitro, MC38-hCD24 showed slightly slower tumor growth in vivo (Fig. S5, D-E). Therefore, mice were inoculated with higher number of MC38-hCD24 cells (1 × 10⁶). In this model, G01 displayed dose-dependent antitumor response, achieving tumor growth inhibition of 62.0% at a low dose of 3 mg/kg and 81.6% at a high dose of 10 mg/kg (Fig. 4, K-L, and Fig. S5, F-G). Importantly, combination of the equimolar cG7 and TGFBR2-Fc only yielded 58.8% inhibition. Importantly, during the treatment, mice remained highly active with steady weight gain (Fig. 4M), indicating again that systemic administration of G01 did not trigger severe toxicity. Furthermore, simultaneous blockade of TGF-β/Smad and CD24/Siglec-10 signaling pathways effectively suppressed tumor proliferation, evidenced by significant reduction in the proportion of Ki-67⁺ cells in G01-treated tumors (Fig. 4, N-O).

Taken together, these data demonstrate the superior anti-tumor efficacy of G01 in multiple mouse tumor models.

### G01 remodels the fibrotic TME and suppresses EMT in vivo by depleting TGF-β1

To understand the underlying mechanisms of G01-mediated anti-tumor efficacy, we first investigated its capability to dismantle the fibrotic stroma barrier of the tumor tissues. We assessed intratumoral collagen deposition, a major component of the fibrotic barrier, in MC38-hCD24 tumor sections. Masson’s trichrome staining revealed that G01 and TGFBR2-Fc treatments significantly diminished the collagen-deposited areas compared to PBS and cG7 treatments (Fig. 5A). Sirius red staining further validated this collagen barrier disruption: tumors treated with G01 exhibited a markedly looser and more sparsely spreaded collagen network. These findings confirm that G01 can alleviate stromal desmoplasia in vivo by inhibiting collagen deposition.

**Fig. 5.**
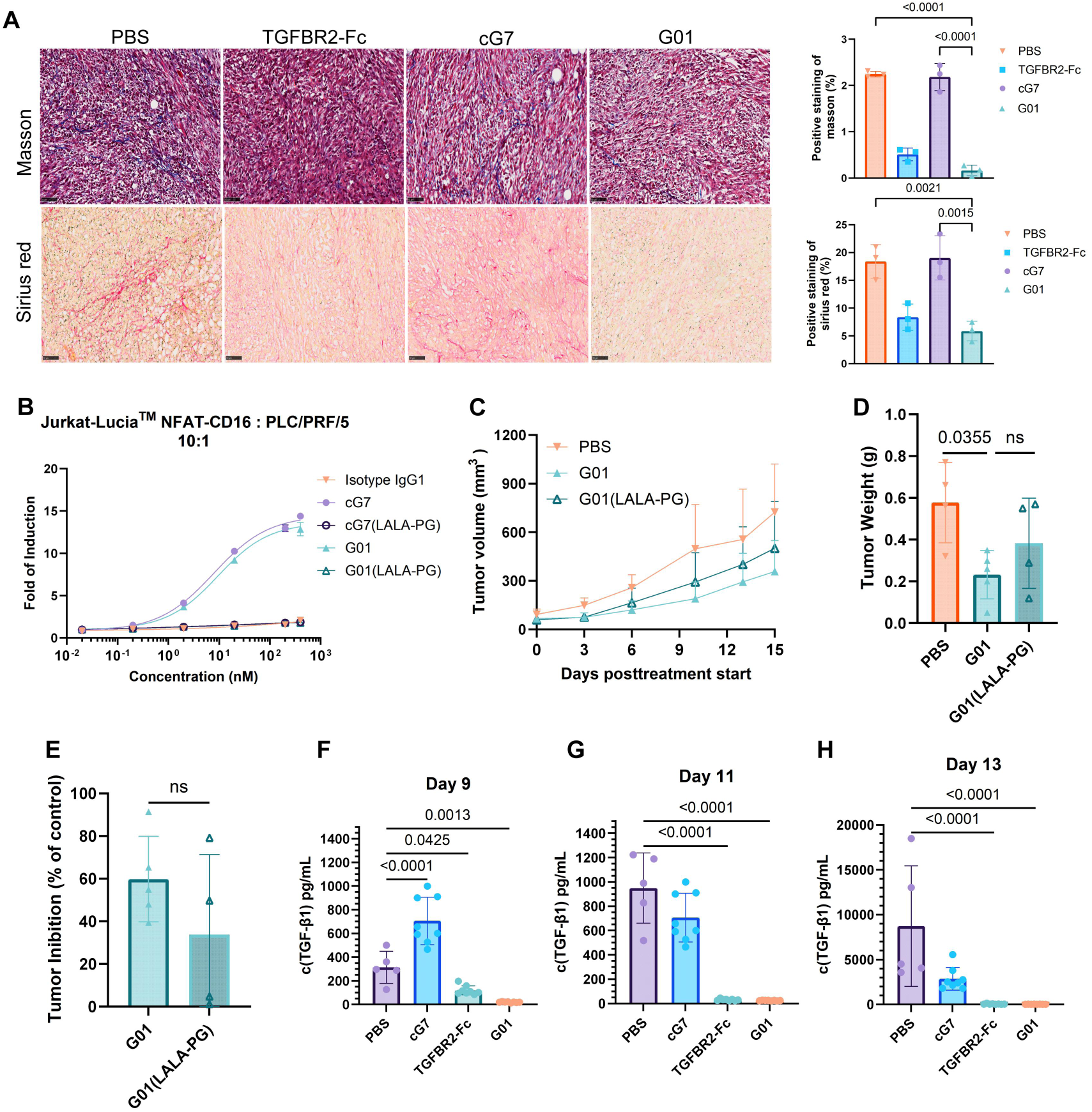
G01 remodels the TME and also leverages Fc-mediated effector functions to suppress tumor progression. **(A)** Representative images and quantitative analysis of intratumoral collagen deposition, assessed by Masson’s trichrome and Sirius Red staining in MC38-hCD24 tumor tissues. Scale bar, 50 μm. Tumors were harvested from C57BL/6J mice treated intraperitoneally for a total of six doses with G01, cG7, TGFBR2-Fc, and PBS. n = 3, and data represent mean ± SD. Statistical significance was determined by one-way ANOVA with Tukey’s post hoc test. **(B)** ADCC evaluation using the CD16a reporter assay. Jurkat-Lucia^TM^ NFAT-CD16 cells were co-cultured with PLC/PRF/5 at an effector-to-target (E:T) ratio of 10:1. The effector-silent variant of cG7 and G01, cG7 (LALAPG) and G01 (LALAPG) were used as controls. **(C to E)** Comparison of the therapeutic efficacy of G01 and G01 (LALAPG) in a humanized xenograft model. NCG mice were reconstituted with human PBMCs via tail vein injection and subcutaneously inoculated with PLC/PRF/5 cells. Mice received intraperitoneal injections of PBS, G01 (10 mg/kg), or G01 (LALAPG) (10 mg/kg) every two days for a total of five doses when tumor volumes reached 50-100 mm³. n = 4 for PBS and G01 (LALAPG) groups, n = 5 for G01 group. (C) Tumor growth kinetics. (D) Terminal tumor weights at day 15 post-treatment. (E) Tumor growth inhibition (TGI) across treatment groups. Data represent mean ± SD, and statistical significance was determined by one-way ANOVA with Tukey’s post hoc test (D) or unpaired two-tailed Student’s t-test (E). **(F to H)** Quantification of plasma TGF-β1 by ELISA at indicated time points after different treatments. n = 5 for PBS, n = 8 for G01, cG7 and TGFBR2-Fc groups. Data represent mean ± SD, and statistical significance was determined by one-way ANOVA with Tukey’s post hoc test.

TGF-β1 is a potent inducer of EMT, driving metastasis. To determine whether G01 treatment could abrogate this process, we examined how downstream EMT markers (N-cadherin and vimentin) and fibroblast activation protein (FAP) changed upon different treatments. Western blot analysis demonstrated that G01 indeed suppressed vimentin expression, potentially contributing to EMT disruption (Fig. S5I-K).

As G01 demonstrates potent effector functions, including ADCC and ADCP, against CD24-expressing tumor cells in vitro, we determined whether these effector functions contribute to the efficacy of G01 in vivo. We first engineered Fc-silenced variants of both G01 and cG7 by introducing the LALAPG mutations into their Fc domains. Using the FcγRIIIa reporter assay, we confirmed that while intact G01 and cG7 robustly activated the ADCC signaling cascade, their respective LALAPG counterparts failed to induce reporter luminescence, validating the abrogation of Fc-effector engagement (Fig. 5B).

Since human IgG1 antibodies exhibit suboptimal affinity for murine Fcγ receptors, immunocompetent mouse models are not suitable to evaluate how effector functions contribute to G01 efficacy in vivo. We therefore employed a humanized mouse model by reconstituting the immune system of severely immunodeficient NCG mice with human PBMCs. While intravenous administration of intact G01 suppressed tumor progression as expected, the LALAPG variant that is eliminated with the effector functions showed inferior tumor growth inhibition (Fig. 5, C-D). Specifically, intact G01 achieved tumor growth inhibition (TGI) of 59.83%, whereas the TGI for the effector-silent variant dropped to 33.77% (Fig. 5E). Though not reaching statistical significance, it should be noted that the high inter-individual variability in PBMC engraftment efficiency, a common phenomenon in immune reconstitution of humanized mouse models, may contribute to the large variation of anti-tumor responses in the same treatment group.

Prompted by the robust clearance of TGF-β1 by G01 in vitro (Fig. 2J-K), we sought to investigate whether G01 could similarly drive the efficient depletion of TGF-β1 in vivo. Mice bearing subcutaneous MC38-hCD24 tumors were treated every 2 days for a total of three doses with G01, or equimolar doses of cG7 or TGFBR2-Fc. Blood was collected at indicated time points for TGF-β1 quantification using ELISA (Fig. S5H). Intriguingly, on day 9 (24 hours after the initial dose), treatment with cG7 resulted in a significant elevation of systemic TGF-β1(Fig. 5F). We postulate that this unexpected increase reflects a compensatory anti-inflammatory response: following the opsonization of tumor cells by cG7, TAMs undergo robust phagocytosis(15), triggering a massive local release of TGF-β1 to orchestrate tissue repair and prevent local hyperinflammation(31–33), which subsequently penetrates into the circulation. Strikingly, although G01 shares the same CD24-binding domain as cG7, it efficienlty depleted TGF-β1 at 24 hours post-treatment. This suggests that G01 leverages hCD24 on tumor cells to degrade TGF-β1 locally within the tumor tissue, thereby effectively cutting off the supply to systemic TGF-β1 elevation. Such robust suppression was sustainable during multiple dosing cycles. On days 11 and 13, plasma TGF-β1 in the G01 and TGFBR2-Fc groups remained significantly lower than that in the PBS cohort, demonstrating that G01 mediates durable clearance of circulating TGF-β1 (Fig. 5G-H).

In summary, these results demonstrate that G01 effectively dismantles the fibrotic shield and suppresses EMT by exhausting TGF-β.

### G01 reprograms the TME to drive immune reactivation

To further understand how G01 reprograms the TME, we performed RNA sequencing on excised MC38-hCD24 tumors. Compared to cG7 or PBS treatment, G01 induced distinct gene expression alterations (Fig. S6, A-C). Gene Ontology (GO) enrichment analysis demonstrated that G01-induced transcriptional changes were heavily enriched in immune-related cascades, including innate immune responses and phagocytosis, suggesting a shift toward more inflammatory TME (Fig. S6D).

To characterize the specific cellular mediators of this inflamed phenotype, we estimated the abundance of 36 distinct tumor-infiltrating immune cell subsets using transcriptomic deconvolution with ImmuCellAI. Consistent with the GO enrichment, tumors from G01-treated mice exhibited enhanced infiltration of multiple T cell lineages, including CD8⁺ T cells, MHC-independent γδ T cells, and memory CD8⁺ T cells. Additionally, augmented infiltration of natural killer (NK) cells and conventional/monocyte-derived dendritic cells, along with a shift toward a higher M1-to-M2 macrophage ratio was observed for innate immune cell subsets in G01-treated mice (Fig. S6, E-F). Although the high intragroup variation inherent to the in vivo models precluded statistical comparison of these immune lineages, the global immune infiltration score was significantly elevated in the G01 group (Fig. S6G), confirming a robust pan-immune reinvigoration by the treatment.

To underpin the mechanisms of this enhanced immune recruitment, we evaluated various immunomodulatory signaling nodes using GSVA. G01 treatment significantly potentiated the activity of STING, IFN-γ, IFN-α, and IL-2-STAT5 signaling pathways compared to the PBS control; among these, STING and IFN signaling induction by G01 was superior to that by cG7 (Fig. S6H). Additionally, transcriptomic analysis confirmed that G01 robustly upregulated the expression of several lymphocyte-recruiting chemokines (e.g., Ccl11, Cxcl9, Cxcl11, Cxcl12, Cxcl16) as well as key cytotoxic effector molecules (Gzma, Gzmb, Prf1, and Ifng) that are essential for CD8⁺ T and NK cell-mediated tumor killing (Fig. S6I).

Taken together, these results indicate that G01 not only drives the recruitment of innate and adaptive immune cells into the TME, but also licenses their anti-tumor cytotoxicity.

### G01 synergizes with PD-1 blockade to boost anti-tumor efficacy

We next examined whether PD-1 blockade can synergize with G01 for enhanced tumor suppression. Mice bearing MC38-hCD24 tumors were treated with PBS, G01, anti-PD-1 antibody, or G01 and anti-PD-1 combination. Indeed, the combination therapy outperformed either G01 or anti-PD-1 monotherapy, which significantly prolonged survival of the tumor-bearing mice (Fig. S7A-B). These findings demonstrate the potential of concurrent targeting of both categories of immune checkpoints.

To interrogate the synergy between G01 and anti-PD-1 antibody, tumor tissues from the PBS, anti-PD-1 monotherapy, and G01 + anti-PD-1 combination groups were collected and subjected to Data-Independent Acquisition (DIA) proteomic profiling. In total, 8,703 high-confidence proteins were identified. Principal component analysis (PCA) revealed spatial separation of the three treatment groups, with the combination therapy inducing the most profound changes in the proteomic landscape (Fig. S8A). Quantitative reproducibility was high across all biological replicates, evidenced by relative standard deviation (RSD) values below 0.3 (Fig. S8B). Hierarchical clustering further demonstrated that while individual variability existed within the anti-PD-1 monotherapy group, the combination therapy maintained a cohesive and distinct proteomic signature (Fig. S8C).

G01 and anti-PD-1 combination elicited more robust proteomic response compared to PD-1monotherapy (Fig. S8, D-E). We identified more DEPs (|FC| ≥1.2, P<0.05) in the combination group relative to the PBS control than in the monotherapy group. Notably, 21 proteins were uniquely altered only in the combination group, suggesting that G01 not only enhances the effects of anti-PD-1 but also triggers an independent regulatory program (Fig. S8, E-F).

Interestingly, the cell-cycle-related transcription factor E2F1 was identified as one of the most significantly downregulated proteins in the combination group (Fig. 6, A-B). Given that E2F1 is a known driver of colorectal cancer proliferation and its protein stability is post-translationally modulated by TGF-β signaling, we investigated its upstream regulators. Notably, we observed significant upregulation of Bmpr2 in the combination group (Fig. 6A). In accordance with the role of Bmpr2 in antagonizing TGF-β-Smad signaling, its upregulation suggests that G01 treatment suppresses the TGF-β axis, thereby destabilizing E2F1 and promoting tumor cell apoptosis.

**Fig. 6.**
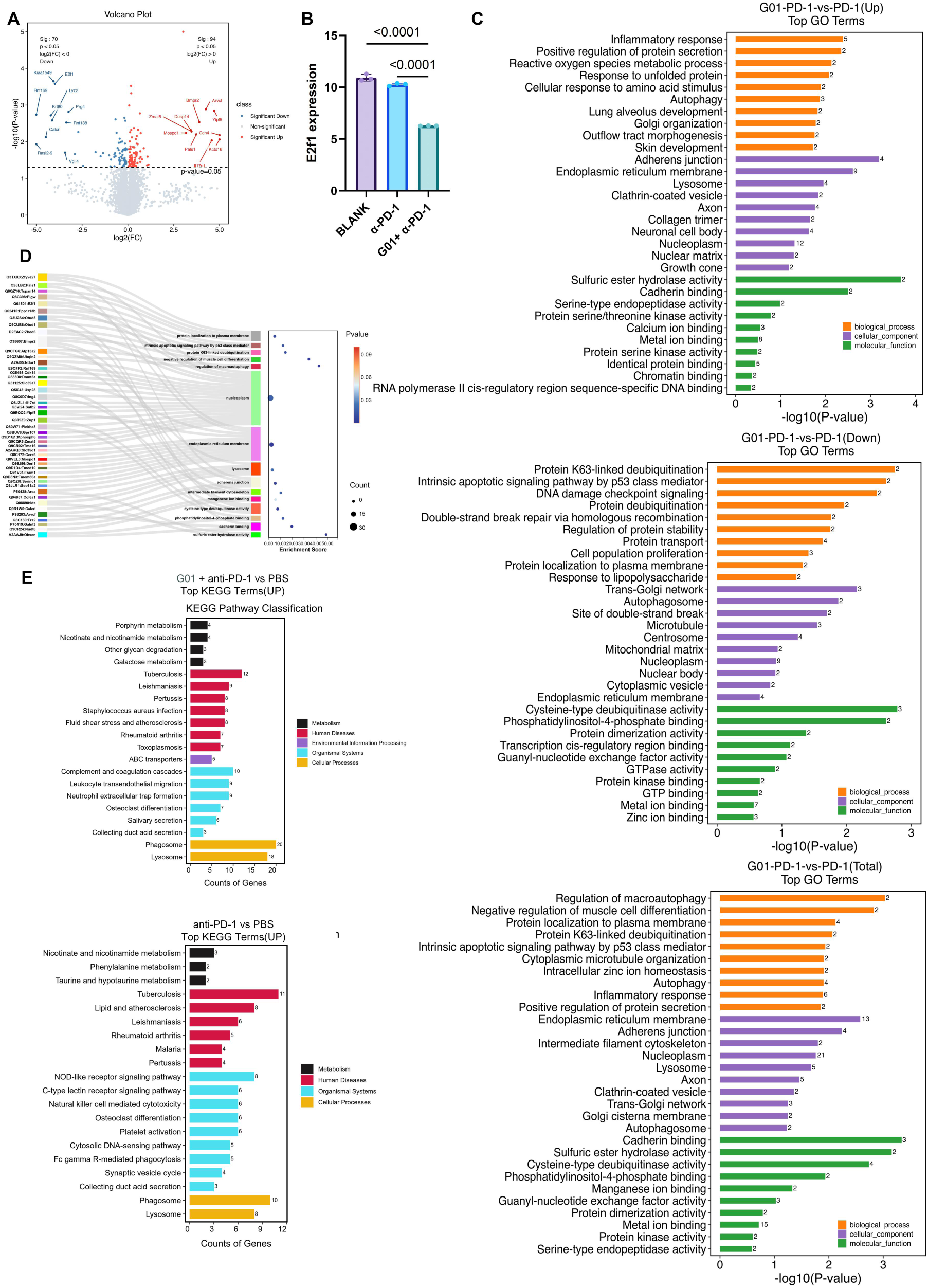
Proteomic profiling reveals that G01 synergizes with PD-1 blockade by modulating numerous inflammatory pathways and suppressing the E2F1 signaling axis. MC38-hCD24 tumor-bearing mice were treated with PBS, anti-PD-1 antibody (1mg/kg), and G01 (10mg/kg) plus anti-PD-1 (1mg/kg) combination (n = 3 per group). Tumors were harvested 6 days after treatment for proteomic analysis. **(A)** Volcano plot illustrating differentially expressed proteins (DEPs) between the G01 + anti-PD-1 combination group and the anti-PD-1 monotherapy group. **(B)** Expression levels of the transcription factor E2F1 across the three experimental groups. Data represent mean ± SD, and statistical significance was determined by one-way ANOVA with Tukey’s post hoc test. **(C and D)** GO enrichment analysis of DEPs in the combination versus monotherapy comparison. (C) Bar chart of enriched GO terms ranked by P value (top to bottom: upregulated, downregulated, and total DEPs). (D) Sankey bubble plot of GO terms (P<0.05). **(E)** KEGG pathway enrichment analysis between PBS group and anti-PD-1 monotherapy group or G01 plus anti-PD-1 combination.

GO and KEGG enrichment analysis revealed that the combination therapy also enriched pathways related to the inflammatory response, macroautophagy, and myoblast differentiation (Fig. 6, C-D). Importantly, the combination group exhibited nearly twofold more upregulated proteins involved in phagosome and lysosome pathways compared to anti-PD-1 monotherapy (Fig. 6E). While E2F1 downregulation typically correlates with reduced autophagy, we observed paradoxical increase in lysosomal and phagosomal proteins. Considering that G01 recognizes a unique epitope on CD24, these data suggest that G01 likely activates a distinct CD24 internalization-dependent program that enhances both tumor cell death and macrophage-mediated antigen presentation, further sensitizing tumors to PD-1 blockade.

### G01 remodels the TME in human PDTFs

While murine models provide valuable mechanistic insights, they often cannot fully recapitulate the heterogeneity and complexity of human solid tumors. To evaluate the translational potential of G01, we employed an ex vivo PDTF platform. This model temporally preserves the native microenvironment of primary human tumors, serving as a high-fidelity surrogate for predicting clinical efficacy.

We first examined CD24 and TGF-β expression in HNSCC. Analysis of single-cell RNA sequencing (scRNA-seq) datasets from two independent cohorts of HNSCC patients revealed that CD24 is specifically overexpressed in malignant epithelial cells, which remains nearly undetectable in most stromal and immune cell populations (Fig. 7, A-F; GSE172577 for A-C, GSE181919 for D-F). On the other hand, TGF-β expression is not restricted to tumor cells, with some immune cell subsets, especially macrophages expressing this immunosuppressive cytokine. Such expression profile supports CD24 targeting with G01 to degrade TGF-β in the TME by preferentially recognizing malignant cells while minimizing the risk of TGF-β-degradation mediated off-tumor toxicity. Importantly, though a subset of plasma cells exhibited moderate co-expression of CD24 and TGFB1, selective targeting of tumor cells but not B cells by G01 (Fig. S1H) suggests that B cells should be spared, therefore minimizing undesired side effects.

**Fig. 7.**
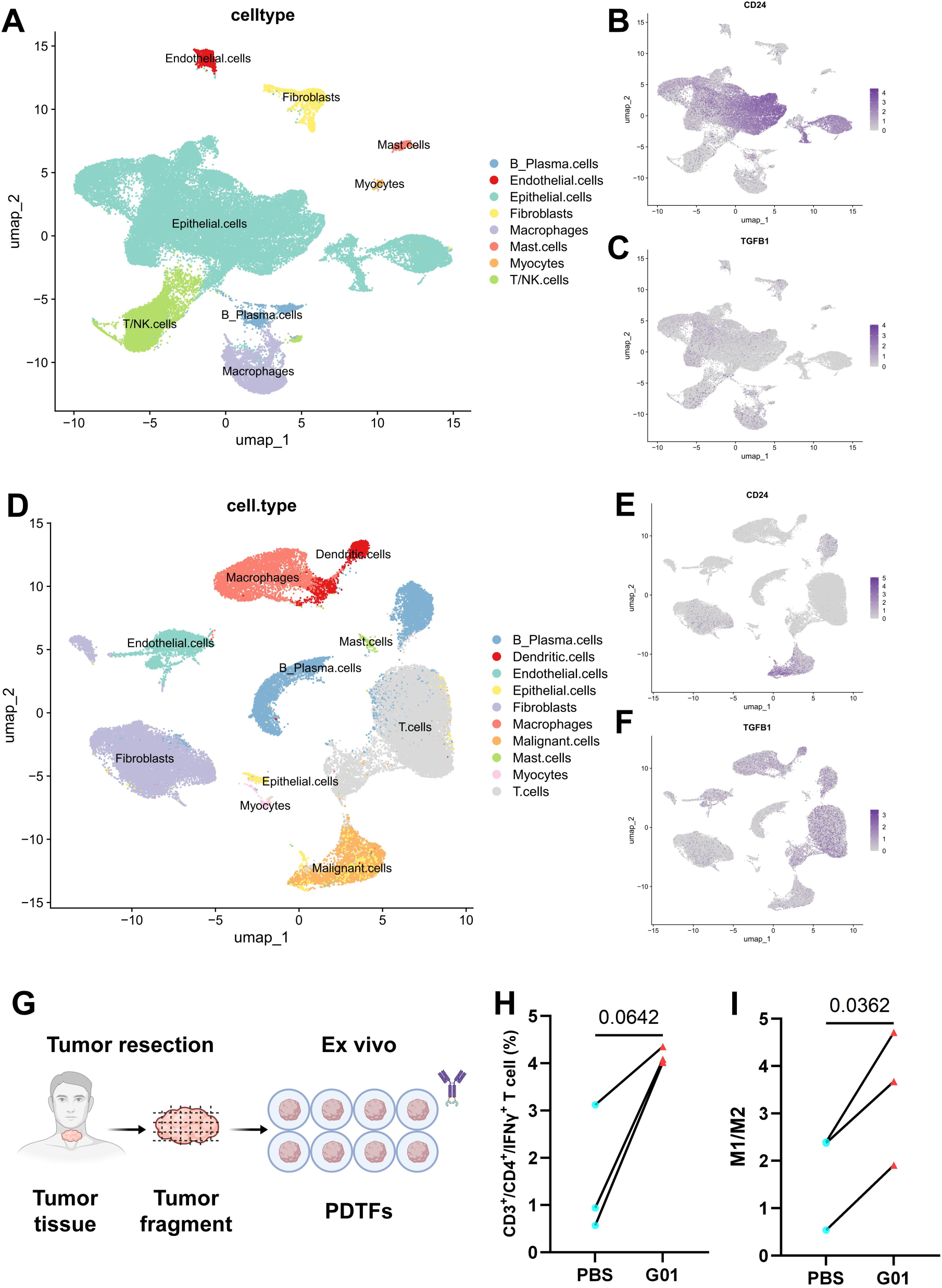
G01 promotes human CD4^+^ T cell activation and M1 macrophage polarization in ex vivo PDTFs. **(A to F)** Uniform Manifold Approximation and Projection (UMAP) of single-cell RNA-sequencing (scRNA-seq) from two independent human head and neck squamous cell carcinoma (HNSCC) cohorts (GSE172577 for A-C; GSE181919 for D-F). Major clusters within the TME annotated according to their characteristic gene markers were shown (A and D), with CD24 (B and E) and TGFB1 (C and F) expression projected onto the respective UMAP plots. **(G to I)** Evaluation of G01-mediated immunomodulation using an ex vivo human PDTF platform. (G) Workflow of PDTF establishment and antibody treatment. Freshly resected human HNSCC tumor fragments (n = 3 independent patients) were cultured ex vivo and treated with either PBS or 100 nM G01 for 48 hours. (H and I) Flow cytometric quantification of immune cells within the PDTFs following treatment. Dot plots showing (H) the percentage of activated CD4^+^ T cells (defined as CD4^+^IFN-γ^+^) in total CD4^+^ T cells and (I) phenotypic shift of TAMs, represented as the ratio of immunostimulatory M1 (CD86^+^) to immunosuppressive M2 (CD206^+^) macrophages. Statistical significance was determined using paired Student’s t-test.

We interrogated the immunomodulatory potential of G01 in the PDTF model. Freshly resected HNSCC fragments (n=3 patients) were treated with G01 (100 nM) or PBS for 48 hours, followed by flow cytometric analysis of the TME (Fig. S9, A-B). Notably, G01 treatment reinvigorated the adaptive anti-tumor immunity, evidenced by the increase of functionally active, IFN-γ^+^ CD4^+^ T cells (Fig. 7H). Additionally, G01 remodeled the innate immune landscape: we observed phenotypic shift in TAMs, characterized by markedly elevated ratio of immunostimulatory M1 (CD86^+^) to immunosuppressive M2 (CD206^+^) macrophages (Fig. 7I). All together, these results suggest that G01 promotes M1 to M2 polarization of TAMs and CD4^+^ T cell activation by disrupting TGF-β signaling and blocking CD24 simultaneously.

In summary, these ex vivo data demonstrate that G01 can effectively overcome TME barriers and synergize innate and adaptive antitumor responses in the human TME, supporting its clinical development.

## DISCUSSION

In this study, we developed the ICTAC platform that harnesses an endocytosing innate immune checkpoint receptor to drive effective internalization and degradation of extracellular proteins. Leveraging the co-dysregulation of CD24 and TGF-β1 in multiple tumor indications, we designed the ICTAC molecule G01, a bifunctional fusion protein composed of anti-CD24 antibody and the ECD of human TGFBR2. We showed that G01 exerts potent antitumor activity through a “trap-and-degrade” mechanism, driven by the rapid internalization of CD24 to deplete TGF-β1. Unlike multiple previously reported extracellular protein degrading platforms, e.g., LYTAC and kineTAC that utilize tumor-specific receptors for degradation, ICTAC harnesses immune-related receptors for such function, thus can be a modular strategy for depletion of other extracellular or membrane proteins by utilizing other innate immune checkpoints. Such unique design also makes concurrent targeting of both the select innate immune checkpoint and the extracellular protein as expected.

Importantly, our ICTAC molecule G01 offers a novel strategy for TGF-β1 targeting, a long-sought objective in oncology. TGF-β1 acts both as a potent immunosuppressive cytokine and an inducer of EMT, thereby fueling tumor immune evasion and metastasis. Previous TGF-β inhibitors all rely on TGF-β blockade with antibodies, antibody-trap fusion proteins, or small molecules, which have not shown clinical benefits, necessitating alternative TGF-β targeting strategies. G01, however, utilizes CD24 as a vehicle on malignant cells to shuttle sequestered TGF-β1 into the lysosomal pathway. Such activity transforms the tumor cells into a molecular sink to deplete the TGF-β reservoir within the TME and provide sustained inhibition of TGF-β signaling compared to conventional antagonists.

Beyond TGF-β depletion, we found that G01 promotes tumor suppression via multiple mechanisms of action. By simultaneously inhibiting CD24-mediated survival signals and TGF-β 1-driven mesenchymal transitions, G01 reduces collagen deposition. This mechanical loosening of the stroma, combined with ADCC and ADCP activities, promotes the influx of anti-tumor immune cells into previously immune-excluded regions.

Regarding safety, despite CD24 expression on B cells but not other normal cells, G01 exhibited negligible binding to healthy human peripheral blood cells in our assays, thanks to the unique epitope of the parental cG7 antibody. This selectivity minimizes the possibility of systemic B cell depletion associated with some previous reported CD24 antibody candidates, providing a wide therapeutic window essential for its potential transition into clinics.

Importantly, we showed that G01 can remodel the human TME. In PDTFs, G01 reinvigorated CD4^+^ T cells and shifted macrophage polarization toward a pro-inflammatory M1-like phenotype. These data further support testing it in clinics.

Certain limitations of our study should be acknowledged. First, while PDTFs preserve the native TME temporally, they lack the systemic immune-tumor circuitry necessary to evaluate other anti-tumor immune components such as memory T cell responses or systemic immune-related adverse events over prolonged periods. Second, the potential impact of the CD24-G01-TGF-β1 complex on homeostatic CD24 recycling dynamics require further elucidation. Third, differences in binding affinities and tissue distribution of CD24 and TGF-β between murine models and humans mean that our in vivo efficacy data must be carefully extrapolated. While the clinical utility of G01 remains to be validated, the novel framework of leveraging cell-surface recycling innate immune checkpoint receptors to create a molecular sink for immunosuppressive factors provides a rationale for developing more potent strategies to overcome therapeutic resistance in solid malignancies.

## List of Supplementary Materials

Present a list of the Supplementary Materials in the following format.

Materials and Methods

**Fig. S1 to S9**

References (*34–36*)

## Acknowledgments

We thank Department of Otorhinolaryngology, The First Affiliated Hospital, Nanjing Medical University, and the patients who participated in this study, for providing human biological samples.

## Author contributions

Z.C., M.P., J.Z. and B.W. designed the study. Y.H. performed bioinformatic analysis. Z.C., M.P. and Y.H. developed methodologies and designed the figures. M.P., Z.C., Y. Z. and M.G. produced and validated all proteins. Z.C. and Y.H. convinced the internalization of surface proteins by FCM. M.P. performed the co-localization of proteins with lysosome by IF. Z.C. and M.G. performed ELISAs of soluble TGF-β1 in vitro and in vivo. M.P. performed reporter gene assay, ELISA, cell transition and migration, cell lysis, ADCC ADCP related experiments and genomic analysis. M.P. and Z.C. performed all experiments in vivo. Z.C. performed proteomic analysis and PDTF related experiments. Z.L. and X.C. provided tumor tissue from human donors. J.Z. and B.W. secured funding for this work. J.Z., B.W. and X.C. supervised the laboratory activities associated with this study. Z.C. and M.P. wrote the initial draft of this manuscript, which was reviewed and edited by J.Z., B.W. and X.C.

## Data and materials availability

All data are available in the main text or the supplementary materials.

## Supplementary Materials

## MATERIALS AND METHODS

### Study Design

The primary objective of this study was to evaluate the therapeutic efficacy, mechanisms of action, and safety profile of G01, a novel dual-targeting (CD24 and TGF-β1) fusion protein. The study design encompassed in vitro mechanistic assays, in vivo syngeneic mouse tumor models, and ex vivo patient-derived tumor fragments (PDTFs) to ensure rigorous translational relevance. For in vivo studies, mice were randomly assigned to treatment and control groups when tumor volumes reached approximately 50-100 mm³. Sample sizes were determined based on expected effect sizes derived from preliminary pilot studies and standard practices in the field, the number of replicates for each experiment is indicated in the figure legends. Endpoints for in vivo studies were prospectively defined, focusing on tumor volume and survival. Tumor measurements and downstream proteomic and flow cytometric analyses were conducted by investigators blinded to the treatment group assignments. The collection and use of human tumor tissues for PDTFs were approved by the Institutional Review Board, and informed consent was obtained from all participating patients.

### Human specimens

All human samples were collected with written informed consent as approved by the Institutional Review Board (IRB) and obtained from Department of Otorhinolaryngology, The First Affiliated Hospital, Nanjing Medical University, Nanjing, China. Human PBMCs were harvested from healthy volunteers by density gradient centrifugation using Lymphocyte-H cell separation medium. HNSCC tumor samples were obtained from HNSCC patients undergoing surgery.

### Mice

All animal experiments were approved by the Institutional Animal Care and Use Committee (IACUC) of China Pharmaceutical University and performed in a Specific Pathogen-Free (SPF) facility. The following mouse strains were used: NCG mice (NOD/ShiLtJGpt-Prkdc^em26Cd52^Il2rg^em26Cd22^/Gpt, Strain NO. T001475, GemPharmatech), Balb/c nude mice (BALB/cNj-Foxn1^nu^/Gpt, Strain NO. D000521, GemPharmatech) and C57BL/6J mice (C57BL/6JGpt, Strain NO. N000013, GemPharmatech).

### Cell lines

All human cell lines, including PLC/PRF/5, BEL-7402, NCI-N87, BxPC-3, HT-29, A549, HuH-7, Jurkat, THP-1, HL-7702, and mouse cell lines, including MC38, were obtained from the Cell Bank of the Chinese Academy of Sciences (Shanghai, China). All cell lines were routinely tested and confirmed to be negative for mycoplasma contamination. Cells were maintained in specific basal media supplemented with 10% fetal bovine serum (FBS) and 1% penicillin-streptomycin according to the American Type Culture Collection (ATCC) guidelines.

Engineered cell lines that overexpress hCD24, mCD24, or hCD16a were generated by lentiviral transduction following with antibiotic selection. Target protein expression was validated using flow cytometry.

### Analysis of TCGA data

CD24 and TGFB1 expression in tumor versus normal tissues was evaluated using The Cancer Genome Atlas (TCGA) database. Gene expression levels were quantified as log_2-_transformed Transcripts Per Million (log_2 [_TPM + 1]) and compared using the non-parametric Wilcoxon rank-sum test, accounting for the non-normal distribution of the transcriptomic data. Expression patterns were categorized into three tiers based on statistical significance and median fold-change: (i) upregulated (median_tumor >_ median_normal a_nd P < 0.05); (ii) downregulated (median_tumor <_ median_normal_ and P < 0.05); and (iii) not significant (P ≥ 0.05). All statistical analyses were conducted using R software (version 4.3.3). Data visualization was performed using ggplot2 (version 4.0.2) and ggpubr packages.

For survival analysis, clinical metadata and log2-transformed transcriptomic data (log_2 [_TPM + 1]) for the TCGA-LIHC and TCGA-BRCA cohorts were retrieved from the UCSC Xena browser. Overall survival (OS), defined as the interval in months from the date of diagnosis to death or last follow-up, served as the primary clinical endpoint. To ensure the robustness of the prognostic analysis and minimize the impact of non-cancer-related perioperative mortality, patients with a follow-up duration of ≤ 0.1 month or those with missing survival status were excluded. Patients were stratified into high or low expression groups based on the median expression levels of CD24 and TGFB1 within each respective cohort. To evaluate the synergistic prognostic impact of these two markers, four distinct patient subgroups were established: *CD24^low^TGFB1^low^, CD24^low^TGFB1^high^, CD24^high^TGFB1^low^, and CD24^high^TGFB1^high^.* Survival probabilities were estimated using Kaplan-Meier analysis, and differences between subgroups, specifically the *CD24*^low^*TGFB1*^low^ versus *CD24*^high^*TGFB1*^high^ populations were assessed using the two-tailed log-rank test. All statistical analysis was performed with the survival (version 3.5-8) and survminer (version 0.5.2) packages.

For immune profiling and single-sample GSEA (ssGSEA), RNA expression data were retrieved for the TCGA-LIHC and TCGA-BRCA cohorts. The expression levels of CD24 and TGFB1 were extracted from each tumor sample, and the samples were classified into a high-expression group and a low-expression group using the median gene expression level as the threshold.

Subsequently, based on the combined expression status of CD24 and TGFB1, the samples were further divided into four subgroups: CD24-low/TGFB1-low, CD24-low/TGFB1-high, CD24-high/TGFB1-low, and CD24-high/TGFB1-high. An in-house curated T cell exhaustion (T_EX)_-associated gene signature was examined as previously defined (*34–36*). This signature comprised 10 key co-inhibitory receptors and exhaustion-related transcription factors: *PDCD1, CTLA4, LAG3, TIGIT, HAVCR2, TOX, EOMES, BATF, ENTPD1,* and *CXCL13*. The overall activity of this transcriptional signature was estimated for each sample using the single-sample gene set enrichment analysis (ssGSEA) algorithm, implemented via the GSVA package (version 1.50.5) in R. Given the non-normal distribution of the enrichment scores, group-wise statistical comparisons specifically between the *CD24*^low^*TGFB1*^low^ and *CD24*^high^*TGFB1*^high^ subgroups were performed using the non-parametric Wilcoxon rank-sum test. Data were visualized using violin plots.

### G01 production

The sequence encoding the ECD of human TGFBR2 was genetically fused to the C-terminus of the cG7 heavy chain via a flexible (G_4S_)_3 l_inker and cloned into the pTT5 mammalian expression vector. This vector was transfected together with the cG7 light chain vector into HEK293F cells (Gibco) using EZ Trans transfection reagent (Life-iLab, Cat# AC04L092) per the manufacturer’s instructions. Culture supernatants were harvested 5 to 7 days post-transfection and clarified by centrifugation. Proteins were captured by Protein A resin (Cytiva) and further purified using Superdex 200 Increase 10/300 GL column (Cytiva) into a custom stabilization buffer (100 mM NaCl, 100 mM glycine, 10 mM citrate, pH 6.0).

### G01 characterization

The purity and monodispersity of the purified proteins were evaluated by analytical SEC using TSKgel G3000SWXL column (Tosoh Bioscience) coupled to Agilent 1260 Infinity II liquid chromatography system. The molecular weight and structural integrity of the purified proteins were assessed by SDS-PAGE under both reducing and non-reducing conditions. G01 Binding kinetics were measured using Octet RED96 (ForteBio). Anti-Human IgG Fc (AHC) biosensors were pre-hydrated for 10 min in assay buffer (PBS containing 0.1% BSA). 100 nM G01, cG7, or TGFBR2-Fc were loaded onto the AHC biosensors to a threshold of approximately 4 nm.

Following a 3-5 min baseline equilibration, the loaded sensors were dipped into wells containing different concentrations of analytes included CD24-mFc (50, 200, and 400 nM) and human TGF-β1, TGF-β2, and TGF-β3 (2.5, 10, and 20 nM) to monitor association (3-5 min). Dissociation was subsequently monitored for 5-10 min. Data were processed using Octet Data Analysis software version 10.0 (ForteBio). Following reference subtraction using a buffer-only negative control, the equilibrium dissociation constant (K_D)_ was calculated using a 1:1 global fitting model. G01 recognition of surface CD24 was examined using FACS. 4 × 10⁵ CHO, CHO-hCD24, CHO-mCD24, PLC/PRF/5, BEL-7402, and Jurkat-CD16a cells were incubated with 5-fold serial dilutions of G01, cG7, or TGFBR2-Fc (ranging from 400 nM to 0.032 nM) at 4°C for 1h, followed by incubation with Alexa Fluor 488 (AF488)-conjugated goat anti-human IgG or goat anti-mouse IgG secondary antibodies (Yeasen Biotechnology) for 1 h at 4°C. Data was acquired using BD FACSCelesta (BD Biosciences) and analyzed using FlowJo software (version 10.8.1).

### G01 concurrent binding to CD24 and TGF-β1

High-binding 96-well microplates (Corning Costar) were coated with 50 μL/well of human TGF-β1 (Novoprotein, cat. no. CA59) at a concentration of 0.5 μg/mL overnight at 4°C. Plates were blocked with 1% BSA in PBS for 2 h at 37°C, followed by adding 100 μL of 4-fold serially diluted G01 (starting from 100 nM to 6.1 μM) for 2 h. 100 μL of 400 nM CD24-mFc was added and incubated for another 2 h, after which 100 μL of HRP-conjugated goat anti-mouse IgG (1:10,000 diluted) was added and incubating for 1 h. After developing with 100 μL of TMB substrate for 10 min, the reaction was stopped by adding 100 μL of 2 M H₂SO₄, and the optical density (OD) was immediately measured using a microplate reader at 450 nm, with 630 nm as a reference wavelength for background subtraction.

Antibody internalization assay 4 × 10⁵ PLC/PRF/5, BEL-7402, A549, HT29, and MC38-hCD24 cells were first incubated with 100 nM of G01, cG7, or isotype control antibody on ice for 1 h to saturate surface CD24 while preventing internalization. For transient exposure, cells were washed twice with ice-cold PBS to remove unbound antibodies before being resuspended in 200 μL of pre-warmed complete culture medium. For continuous incubation, cells were directly resuspended without washing. Samples were then incubated at 37°C for different times (0, 5, 15, 30, 45, 60, 90, and 120 min), whereas the isotype control samples were maintained at 4°C for 120 min. Antibody internalization was stopped by immediately transferring the samples onto ice, and surface-retained antibodies that are not internalized were detected using AF488-conjugated goat anti-human IgG secondary antibody (1:200 dilution; Yeasen) at 4°C for 1 h. The internalization rate at each time point (t) was calculated based on the decay of surface MFI using the following formula: Internalization (%) = [1 − (MFI_t −_ MFI_background)_ / (MFI_0min −_ MFI_background)_] × 100%, where MFI background represents the fluorescence of control cells stained with the secondary antibody.

### G01/TGF-β1 complex internalization

Equal volumes (50 μL) of 10 μg/mL His-TGF-β1 (Novoprotein) and 30 μg/mL antibody were gently mixed and pre-incubated on ice for 30 min. 4 × 10⁵ MC38-hCD24 cells were then resuspended in the pre-assembled mixtures and incubated on ice for 30 min to saturate surface binding while preventing endocytosis. After washing with ice-cold PBS twice, cells were resuspended in pre-warmed complete DMEM medium and incubated at 37°C for different times (0, 0.5, and 2 h), after which internalization of the complex was stopped by incubating the samples on ice. Cells were co-stained with AF488-conjugated goat anti-human IgG (1:200; Yeasen) and APC-conjugated anti-His antibody (1:20; Yeasen) for 30 min to detect the surface-retained antibody and the TGF-β1 ligand, respectively. AF488 and APC fluorescent signal decay relative to the 0-h baseline was utilized to quantify the complex internalization.

### Confocal microscopy and intracellular trafficking analysis

To verify G01 entry into the acidic intracellular compartments (e.g., endosomes and lysosomes), G01, cG7, and TGFBR2-Fc were first conjugated with the pH-sensitive dye Protonex™ Red 600 SE (AAT Bioquest) at room temperature for 1 h, following with centrifugal ultrafiltration (50 kDa MWCO; Mllipore) to remove excess dye. 5 × 10⁵ PLC/PRF/5 and BEL7402 cells were seeded into glass-bottom imaging dishes and cultured for 24 h. Cells were then incubated with 100 nM of the Protonex-conjugated proteins at 4°C for 10 min to synchronize endocytosis, followed by shifting to 37°C for 2 h to induce internalization. After the incubation, nuclei were counterstained with Hoechst 33342 (1:1000 in PBS; Yeasen) for 3 min at room temperature, and cells were imaged using Nikon Ts2R wide-field inverted fluorescence microscope.

To further evaluate the intracellular trafficking of G01, 5 × 10^5^ PLC/PRF/5 cells were seeded in confocal-grade glass-bottom dishes and cultured for 24 h. Cells were then incubated with 100 nM G01 at 4°C for 30 min, after which they were either fixed immediately (0 h) or shifted to 37° C for 0.5 h and 4 h and fixed. Upon permeabilization with 0.2% Triton X-100, cells were incubated with mouse anti-human Lamp-1 primary antibody (Yeasen) for 1 h at RT, followed by incubation with Alexa Fluor 488-conjugated goat anti-human IgG (1:50, Yeasen) and Cy3-conjugated goat anti-mouse IgG (1:50, Yeasen) for 45 min at RT for G01 and Lamp-1 detection, respectively. Nuclei were counterstained with Hoechst 33342. Images were acquired using Leica STELLARIS 5 confocal microscope (Leica Microsystems) equipped with a 400 × objective and analyzed using the Leica Application Suite X (LAS X) software.

### In vitro TGF-β1 degradation

HEK293T-hCD24, PLC/PRF/5, HT29 cells were seeded in 24-well plates at a density of 4 × 10^5^ cells per well overnight, followed by incubation with 10 nM His-tagged human TGF-β1(Sino Biological), 10 nM TGF-β1 and 20 nM G01, or 10 nM TGF-β1 and 100 nM G01 for 12 and 24 h. Supernatants were harvested, clarified to remove cellular debris, and coated on High-Binding microplates. The plates were incubated with mouse anti-His primary antibody (1:150, T505, SignalWay Antibody) for 2 h, followed by incubation with HRP-conjugated goat anti-mouse secondary antibody (Yeasen) for 1 h. The absorbance was measured at 450 nm and 630 nm using a microplate reader. The percentage of remaining TGF-β1-His was calculated as follows: TGF-β 1 remaining (%) = [ OD_450-630(_t) - OD_450-630(_Blank)] / [ OD_450-630(_TGF-β1) - OD_450-630(_Blank)] * 100%.

### TGF-β/Smad signaling reporter assay

HEK293T/SBE-luc reporter cells (Nanjing Leads Biolabs) were seeded into 96-well plates at a density of 5 × 10^4^ cells per well in 100 μL overnight. 10 ng/mL human TGF-β1 (PeproTech) was premixed with varying concentrations of G01, cG7, or TGFBR2-Fc (ranging from 0.02 nM to 100 nM), added to the culture, and incubated for additional 16 h. Following incubation, cells were lysed using 100 μL of lysis buffer (Yeasen) per well for 5 min on ice, and 20 μL of the lysate was transferred into a white 96-well plate and mixed with 100 μL of Firefly Luciferase assay reagent (Yeasen). Luminescence was measured using a Synergy microplate reader (BioTek Instruments), and the percentage of TGF-β-induced signal was calculated using the following formula: TGF-β induced signal (%) = [(Experimental RLU − Blank RLU) / (Positive control RLU − Blank RLU)] × 100. Half-maximal inhibitory concentration (IC_50)_ was determined using a four-parameter logistic (4PL) regression model in GraphPad Prism version 10.2.3 (GraphPad Software).

### SEB-stimulated IFN-γ release assay

PBMCs were seeded into 96-well culture plates at a density of 2.5 × 10^4^ cells per well. 20 ng/mL TGF-β1, 200 nM of the protein drugs (G01, cG7, TGFBR2-Fc, or Isotype IgG1), and 10 ng/mL SEB were added in the next day and incubated for 72 h. Culture supernatant was harvested and diluted 10-fold for Interferon-gamma (IFN-γ) quantification using the ELISA MAX™ Standard Set for Human IFN-γ (BioLegend) following the manufacturer’s instructions.

### TGF-β1-induced EMT in vitro

PLC/PRF/5 cells were seeded into 6-well culture plates at a density of 5 × 10^5^ cells per well overnight. Following serum starvation with 2% FBS for 12 h, cells were treated with 10 ng/mL TGF-β1 together with 100 nM protein drugs for 72 h. EMT characteristic phenotypic changes, i.e., the transition from a cobblestone-like epithelial monolayer to a scattered, spindle-shaped mesenchymal morphology, were assessed using a Nikon Ts2R inverted microscope.

### In vitro wound healing assay

EMT-6-hCD24 cells were seeded into 6-well plates at a density of 5 × 10^5^ cells per well to reach 100% confluency overnight. Linear scratches were gently introduced across the center of each well using a sterile 200 μL pipette tip. Pre-drawn parallel lines on the plate bottom were used as visual guides to ensure perpendicular scratch and consistent image orientation. Following scratch, cellular debris and detached cells were removed by gently washing the wells. Cells were treated with 10 ng/mL TGF-β1 together with 100 nM of G01, cG7, or TGFBR2-Fc, and wound closure was monitored at 0, 12, and 24 h post-treatment using a Nikon Ts2R inverted microscope. For rigorous quantification, two representative fields of view were captured per well, and the cell-free wound area was measure using ImageJ software (NIH). The cell migration rate was calculated using the following formula: Migration rate (%) = [(Initial wound area at 0 h − Wound area at time t) / Initial wound area at 0 h] × 100.

### Transwell invasion assay

The invasive capability of CD24^+^ tumor cells was evaluated using Matrigel-coated Transwell chambers (8.0-μm pore size; Millipore). The upper surface of the porous membrane was pre-coated with 20 μl of Matrigel (BD Biosciences) diluted 1:2 in basal DMEM and allowed to polymerize for 1 h at 37°C. BEL-7402 and Huh-7 cells were seeded into the upper chambers at a density of 5 × 10^4^ cells per well and treated with 200nM cG7, G01, or TGFBR2-Fc. The lower chambers were filled with DMEM supplemented with 5% FBS. After 36-h incubation, non-invading cells remaining on the upper surface of the membrane were mechanically removed. Cells that had successfully invaded into the lower surface of the membrane were fixed, stained with 0.1% crystal violet, and imaged. Four random fields of view per insert were selected.

### ADCP analysis

Primary macrophages were differentiated from PBMCs by seeding into 6-well plates overnight. Non-adherent cells were gently removed by PBS washes, and adherent cells (monocytes) were cultured in RPMI 1640 supplemented with 50 ng/mL M-CSF (ABclonal), with half of the medium replaced every 2 days. Cells were harvested between days 7 and 9 upon exhibiting typical macrophage adherence and morphology. For M2 macrophage polarization, 50 ng/mL TGF-β1 (PeproTech) and 50 ng/mL IL-10 (PeproTech) were added to the M-CSF-containing culture on day 3 or 4, and fully polarized M2 macrophages were similarly harvested between days 7 and 9. THP-1-derived macrophages were differentiated by first seeding the cells in non-tissue culture-treated 6-well plates and stimulated with 100 ng/mL phorbol 12-myristate 13-acetate (PMA; Yeasen) for 24 h. M1 macrophages were further polarized by incubation with 20 ng/mL IFN-γ (Meilunbio) and 100 ng/mL LPS (MedChemExpress) for 24 h, while M2 macrophages were induced with 20 ng/mL IL-4 (PeproTech) and 20 ng/mL IL-13 (ABclonal) for 24 h. The phenotypes of the differentiated macrophages were examined by FACS using the following fluorophore-conjugated antibodies: APC anti-CD11b (clone M1/70, BioLegend), BV421 anti-human CD206 (clone 15-2, BioLegend), and PE anti-human Siglec-10 (clone 5G6, BioLegend). For ADCP analysis, 4 × 10^5^ BEL-7402 cells were first labeled with 1 μM Calcein-AM (Yeasen) for 30 min at 37°C, and incubated with 2 × 10^5^ PBMC-derived M2 macrophages (effector-to-target (E:T) ratio of 1:2) and 100 nM of G01, cG7, or an isotype control IgG1 for 4 h at 37°C. Cells were stained with APC anti-human CD11b antibody for 45 min at 4°C, and the phagocytosis rate was calculated as the percentage of Calcein and CD11b double-positive cells within the total viable CD11b-positive macrophages.

### ADCC analysis

ADCC was evaluated using a luciferase-based reporter assay. MC38-hCD24 cells were seeded into 96-well culture plates at a density of 2 × 10^4^ cells per well and incubated with 50 μL of antibodies (6-point, 10-fold serial dilutions, with a maximum concentration of 200 nM) for 1 h. Jurkat-Lucia NFAT-CD16 reporter cells (InvivoGen) were used as effector cells, and 2 × 10^5^ cells were added to each well for an effector-to-target (E:T) ratio of 10:1 and incubated for 6 h. After incubation. 80 μL supernatant was transferred to a white opaque 96-well plate, mixed with 80 μL of Renilla luciferase detection reagent, and incubated for 10 min. Luminescence was quantified using a BioTek Synergy microplate reader, and ADCC activity was evaluated by calculating the induction fold using the following formula: Fold of induction = (induced RLU − background RLU) / (no antibody control RLU − background RLU).

In vitro cell viability and growth inhibition assay The anti-proliferative effect of G01, cG7, and TGFBR2-Fc was evaluated using MTT assay. HL-7702, BEL-7402, and HuH-7 cells were seeded into 96-well plates and treated with a six-point concentration gradient (0.02, 0.2, 2, 20, 100, and 200 nM) of G01, cG7, or TGFBR2-Fc. After 72 h, 20 μL of MTT solution (5 mg/mL in PBS) was added to each well, and the plates were incubated for 4 h. The supernatant containing unreacted MTT was removed, and the resulting intracellular formazan crystals were solubilized by adding 150 μL of DMSO per well. The absorbance was measured at 570 nm using a microplate reader, and viability was calculated as a percentage relative to the untreated control.

### Tumorigenicity validation

To confirm that eGFP-Luciferase and hCD24 overexpression did not alter the biological fitness of the parental cell lines, in vitro proliferation and in vivo tumorigenicity were evaluated. For in vitro assessment, wild-type (WT) and engineered cells (PLC/PRF/5-Luc vs. WT PLC/PRF/5; MC38-hCD24 vs. WT MC38) were seeded into 96-well plates at a density of 2 × 10^3^ cells per well. Every 24 h over a 7-day period, 10 μL of Cell Counting Kit-8 (CCK-8) reagent was added to the plate, followed by a 4-h incubation. Absorbance was measured using a microplate reader at a primary wavelength of 490 nm with a reference wavelength of 630 nm. Net absorbance (OD490-OD630) was calculated to examine cellular proliferation.

To verify that hCD24 overexpression did not compromise tumorigenicity, female C57BL/6J mice were subcutaneously inoculated with 5 × 10^5^ or 1 × 10^6^ WT MC38 or MC38-hCD24 cells. Tumor growth kinetics was monitored and compared.

### Tumor treatment studies

For all in vivo efficacy studies, antibodies were administered via intraperitoneal (i.p.) injection every two or three days. When tumor volumes reached 50-100 mm3, mice were randomly assigned to different treatment groups (n = 4-8 mice per group) including: high-dose G01 (10 mg/kg), low-dose G01 (3 mg/kg), equimolar cG7 (8.3 mg/kg), equimolar TGFBR2-Fc (4.6 mg/kg), combination of cG7 and TGFBR2-Fc, and vehicle control (100 μL PBS). Throughout the study, body weights of the mice were continuously monitored to evaluate systemic toxicity.

For orthotopic liver tumor model, male Balb/c nude mice were inoculated with 2 × 10^6^ PLC/PRF/5-Luc cells into the hepatic lobe. Tumor progression was monitored on day 0, 7, and 17 post treatment (corresponding to day 7, 14 and 24 after transplantation) by bioluminescence imaging. Mice were euthanized on day 17 for endpoint analysis.

For subcutaneous models, 1 × 10^6^ BEL-7402 or MC38-hCD24 cells were inoculated into the right flank of male Balb/c nude mice or female C57BL/6J mice, respectively. Tumor volumes were measured as V=0.5 × length × width^2^ using digital calipers every 2 days for BEL-7402 model and every 3 days for MC38-hCD24 model. At the pre-defined study endpoint (day 16 for BEL-7402 model and day 13 for MC38-hCD24 model), mice were euthanized, and tumors were completely excised and weighed. The tumor growth inhibition (TGI) was calculated using the formula: TGI (%) = [1 - (Tumor weight of treatment group / Tumor weight of vehicle group)] × 100%. For BEL-7402 model, Kaplan-Meier survival curves were also generated. Tumor volumes did not exceed 2000mm3 at any time during the studies.

### Histopathological and IHC analysis

At the study endpoint, tumors and kidneys from the BEL-7402 cohort were harvested and fixed with 4% paraformaldehyde. Tissues were then paraffin-embedded and sectioned (Pathology Platform, China Pharmaceutical University). For toxicity evaluation, kidney sections were stained with hematoxylin and eosin (H&E). Renal edema was assessed using tubular epithelial cell swelling and luminal narrowing. For tumor proliferation assessment, tumor sections were stained for Ki67 by Pathology Platform, China Pharmaceutical University. Images were acquired using an S60 whole-slide scanner, and the percentage of Ki67-positive tumor cells were quantified by randomly selecting three to five high-power fields per slide. Tumors of MC38-hCD24 model were processed similarly.

### Tumor extracellular matrix remodeling and fibrosis

Tumors were collected from the MC38-hCD24 cohort at the endpoint, fixed, paraffin-embedded, sectioned, and examined using standard Masson’s trichrome staining (Pathology Platform, China Pharmaceutical University) to examine collagenous extracellular matrix depostion. Tumor sections were also stained with Sirius Red to check collagen I fibers. Fibrosis was quantified as the percentage of the positively stained area relative to the total tumor area using ImageJ.

### Analysis of cancer-associated fibroblasts and EMT

MC38-hCD24 tumors were collected at the study endpoint, rinsed with ice-cold PBS, fractionated into ∼50 mg aliquots, and snap-frozen in liquid nitrogen to preserve protein integrity. Tumor fragments were thawed, immersed in 300-500 μL of RIPA lysis buffer supplemented with 1× protease and phosphatase inhibitor cocktails, and homogenized using a 3D cryogenic tissue pulverizer at 60 Hz for 4 cycles of 30 seconds each, interspaced with 30-second cooling intervals. Then ultrasonic homogenization was performed using a probe sonicator on ice to prevent protein denaturation. Homogenates were subsequently centrifuged at 12,000 rpm for 20 min at 4°C. Supernatants were collected, and total protein concentrations were quantified using a BCA protein assay kit. The lysates were resolved by SDS-PAGE, followed by membrane transfer and blot with the following antibodies: mouse anti-FAP (Abmart, 1:1,000), rabbit anti-N-Cadherin (HUABIO, 1:5,000), rabbit anti-vimentin (Cell Signaling, 1:1,000). Mouse anti-GAPDH (Proteintech, 1:100,000) was used as an internal loading control. Membranes were incubated with HRP-conjugated goat anti-mouse or anti-rabbit IgG secondary antibodies (Yeasen, 1:10,000) and developed using enhanced chemiluminescence. To facilitate quantitative comparison of samples across multiple gels, a positive control sample was loaded onto every gel as an inter-gel calibrator. Target protein band intensities were first normalized to the internal loading control (GAPDH) within the same lane. Subsequently, these intra-lane normalized values were divided by the normalized value of the inter-gel calibrator from the respective gel to assure accuracy of cross-gel statistical comparisons.

### ADCC in vivo using humanized mouse tumor models

To investigate whether ADCC contributes to G01-mediated tumor regression in vivo, the antitumor efficacy of wild-type G01 and its Fc-silenced mutant (G01-LALAPG) was compared using an immune-reconstituted mouse model. Five-week-old male NCG mice were subcutaneously inoculated with 2 × 10^6^ PLC/PRF/5-luc cells. One week post tumor implantation, human immune reconstitution was performed by intravenously injecting 1 × 10^7^ human PBMCs (in 150 μL PBS) via the tail vein. Two weeks following PBMC engraftment, blood was collected retro-orbitally to evaluate human leukocyte chimerism via flow cytometry. Mice exhibiting successful immune reconstitution, defined as the proportion of human CD45^+^ cells constituting more than 20% of total CD45^+^ cells [hCD45^+^ / (hCD45^+^ + mCD45^+^) ≥ 20%], and bearing tumor volumes of 50 to 100 mm3 were randomized and treated by intraperitoneal (i.p.) injections of PBS (100 μL), wild-type G01 (10 mg/kg), or G01-LALAPG mutant (10 mg/kg) every two days for a total of 7 doses. Mice were euthanized 15 days post treatment, and serum and tumor tissues were harvested for downstream analysis.

### Quantitation of TGF-β1 in mouse plasma

Whole blood samples were collected at designated time points into tubes containing sodium EDTA as an anticoagulant. Blood was centrifuged at 2,000 × g for 2 min at 4℃. The resulting supernatant was carefully transferred to new tubes and subjected to centrifugation at 10,000 × g for 10 min at 4℃ to eliminate residual platelets. The supernatant was collected for analysis. The concentration of TGF-β1 in the plasma was quantified using a commercial TGF-β1 ELISA kit (Invitrogen, Cat# 88-8350-22) according to the manufacturer’s instructions. OD was measured using a microplate reader at an absorbance wavelength of 450 nm, with a reference wavelength of 630 nm for background subtraction.

### RNA sequencing and transcriptomic analysis

MC38-hCD24 tumors were collected (n = 3 per group) at the study endpoint. Approximately 50 mg of tumor tissues was isolated, washed with ice-cold RNase-free PBS and snap-frozen in liquid nitrogen. RNA isolation, library preparation, and sequencing were performed by LC-Bio Technologies (Hangzhou, China). Raw sequencing data were aligned to the mouse reference genome (mm10), and gene expression was normalized and expressed as log_2(_TPM + 1).

Differential expression analysis between the treatment groups and the PBS control group was performed using the DESeq2 package (version 1.14.1) in the R software environment (version 3.6.3). Hierarchical clustering of differentially expressed genes (DEGs) was visualized using the heatmap R package (version 1.0.12). The identified DEGs were subjected to enrichment analysis for GO terms and Kyoto Encyclopedia of Genes and Genomes (KEGG) pathways. To quantitatively evaluate immune cell infiltration within the TME, single-sample gene set enrichment analysis (ssGSEA) was performed. Normalized gene expression profiles were analyzed using the ImmuCellAI-mouse tool, which estimates the relative abundances of 36 immune cell types, including 13 T cell subsets and 15 other immune cells (including B cells, NK cells, monocytes, macrophages, neutrophils, and dendritic cells), and calculates an overall immune infiltration score for each sample.

### Proteomics

Tumor tissues from the MC38-hCD24 model were collected one week post treatment and snap-frozen in liquid nitrogen. Total proteins were extracted and quantified, and sample concentrations were normalized across all groups using lysis buffer. Dithiothreitol (DTT) was then added to a final concentration of 5 mM, and the mixture was incubated at 55°C for 30 min. After cooling to room temperature, samples were alkylated with 10 mM iodoacetamide (IAA) for 15 min at room temperature, followed by acetone precipitation. The resulting pellets were air-dried for 2-3 min and resuspended in 100 μL of 50 mM NH₄HCO₃. Proteins were digested using Trypsin-TPCK (1 mg/mL) at an enzyme-to-substrate ratio of 1:50 (w/w) overnight at 37°C.

Digested peptides were lyophilized and desalted using SOLA™ SPE 96-well plates. Briefly, cartridges were sequentially activated with 200 μL of methanol (three times) and equilibrated with 200 μL of 0.1% formic acid (FA) in water (three times). Samples (50-500 μL) were loaded under a controlled vacuum flow rate of ∼1 mL/min (approx. 1 drop/sec) and reloaded once to maximize binding efficiency. After washing with 200 μL of 0.1% FA in water (three times), peptides were eluted with 150 μL of 50% acetonitrile (ACN) containing 0.1% FA (three times, yielding 450 μL total eluate) and vacuum-dried. Prior to MS analysis, indexed Retention Time (iRT) standards were spiked into each sample at a 1:20 (v/v) ratio for retention time calibration. Equal amounts of peptides from each sample were subjected to liquid chromatography separation using an EASY-nLC 1200 system (Thermo Fisher Scientific). The mobile phases consisted of 0.1% FA in water (buffer A) and 0.1% FA in ACN (buffer B). Separation was achieved using a 30-min preprogrammed gradient: 5-22% B for 20 min, 22-37% B for 4 min, 37-80% B for 3 min, and maintained at 80% B for 3 min. The LC system was interfaced with a timsTOF Pro mass spectrometer (Bruker). MS parameters were configured as follows: capillary voltage, 1.4 kV; dry gas temperature, 180°C; dry gas flow rate, 3.0 L/min. DIA was performed with a m/z scan range of 100 to 1700, an ion mobility range of 0.7 to 1.3 Vs/cm², and a collision energy ranging from 20 to 59 eV. Raw DIA MS data were processed using Spectronaut Pulsar™ 18.4 (Biognosys, Switzerland). The search engine parameters included a maximum of two missed cleavages, with carbamidomethylation of cysteine (C) set as a fixed modification, and methionine oxidation (M) and N-terminal acetylation set as variable modifications. A 1% false discovery rate (FDR, q-value < 0.01) was applied at both the precursor and protein levels. To ensure high data fidelity, proteins were filtered to retain those with ≥ 1 unique peptide. Valid quantitative values were required in ≥ 2 samples across all cohorts, or presence in ≥ 50% of the replicates within at least one experimental group. Missing values in qualifying groups were imputed using the intra-group mean, whereas remaining missing values were replaced with half of the matrix minimum. Data were subsequently median-normalized and log2-transformed. Data reproducibility and intra-group consistency were confirmed by evaluating the Relative Standard Deviation (RSD) of biological replicates (n = 3 per group), which yielded values < 0.3 across all cohorts.

Furthermore, global sample relationships were visualized using PCA and Complete-link hierarchical clustering based on Euclidean distance. DEPs were defined using a threshold of fold change (FC) ≥ 1.2 (significantly upregulated) or ≤ 0.83 (significantly downregulated) coupled with a P-value < 0.05. Bi-directional hierarchical clustering with Z-score standardization was utilized to generate heatmaps illustrating the expression patterns. For functional interpretation, DEPs were annotated using the GO database across Biological Process (BP), Cellular Component (CC), and Molecular Function (MF) categories. Pathway analysis was conducted utilizing the Kyoto Encyclopedia of Genes and Genomes (KEGG) database. Furthermore, protein-protein interaction (PPI) networks among the DEPs were constructed using the STRING database to elucidate coordinated functional mechanisms.

### Generation of PDTFs

Freshly collected HNSCC tumor tissues were cut into 1-2 mm³ fragments on ice, and fragments from distinct regions of the tumor were pooled together for ex vivo culture. An engineered ECM hydrogel consisting of 4 mg/ml phenol red-free high-concentration Matrigel (Yeasen), 1 mg/ml rat-tail Collagen I (Sigma-Aldrich), and 1.1% sodium bicarbonate (to neutralize pH and initiate collagen fibrillation) was formulated on ice using basal culture medium consisting of high-glucose DMEM (Keygen) supplemented with 10% FBS, 1 mM sodium pyruvate, 2 mM L-glutamine, 1× MEM Non-Essential Amino Acids (Sigma-Aldrich), and 1% penicillin-streptomycin. 96-well plates were then pre-coated with 40 μl of the ECM mixture per well and incubated at 37°C for 20-30 min for ECM solidification. PDTFs were washed extensively with basal tumor medium, and placed onto the solidified base layer. Subsequently, PDTFs were overlaid with an additional 40 μl of the ECM mixture and incubated at 37°C for another 20-30 min to ensure complete encapsulation. 120 μl of basal tumor medium was added to each well.

### PDTF treatment

PDTFs were treated with 100 nM G01 or an equal volume of vehicle control (PBS) and maintained at 37°C for 48 h. At this time, PDTFs were collected and digested in serum-free RPMI 1640 medium supplemented with 1 mg/mL collagenase IV and 20 U/mL DNase I for 1 h at 37°C with continuous agitation. The resulting mixture was filtered through 70-μm cell strainers to generate single-cell suspensions, then surface-stained for CD45-FITC, CD3-APC, CD4-BV510, CD8-PE-Cy7, CD206-BV421, and CD86-APC (Biolegend). Cells were then fixed and permeabilized using the Foxp3 / Transcription Factor Staining Buffer Set (eBioscience) according to the manufacturer’s instructions, and intracellularly stained with IFN-γ-PE and CD68-PE (eBioscience). Samples were acquired on a BD FACS Celesta (BD Biosciences) and analyzed using FlowJo software v10.8.1 (TreeStar Inc.).

### Fig. S1 to S9

**Fig S1.**
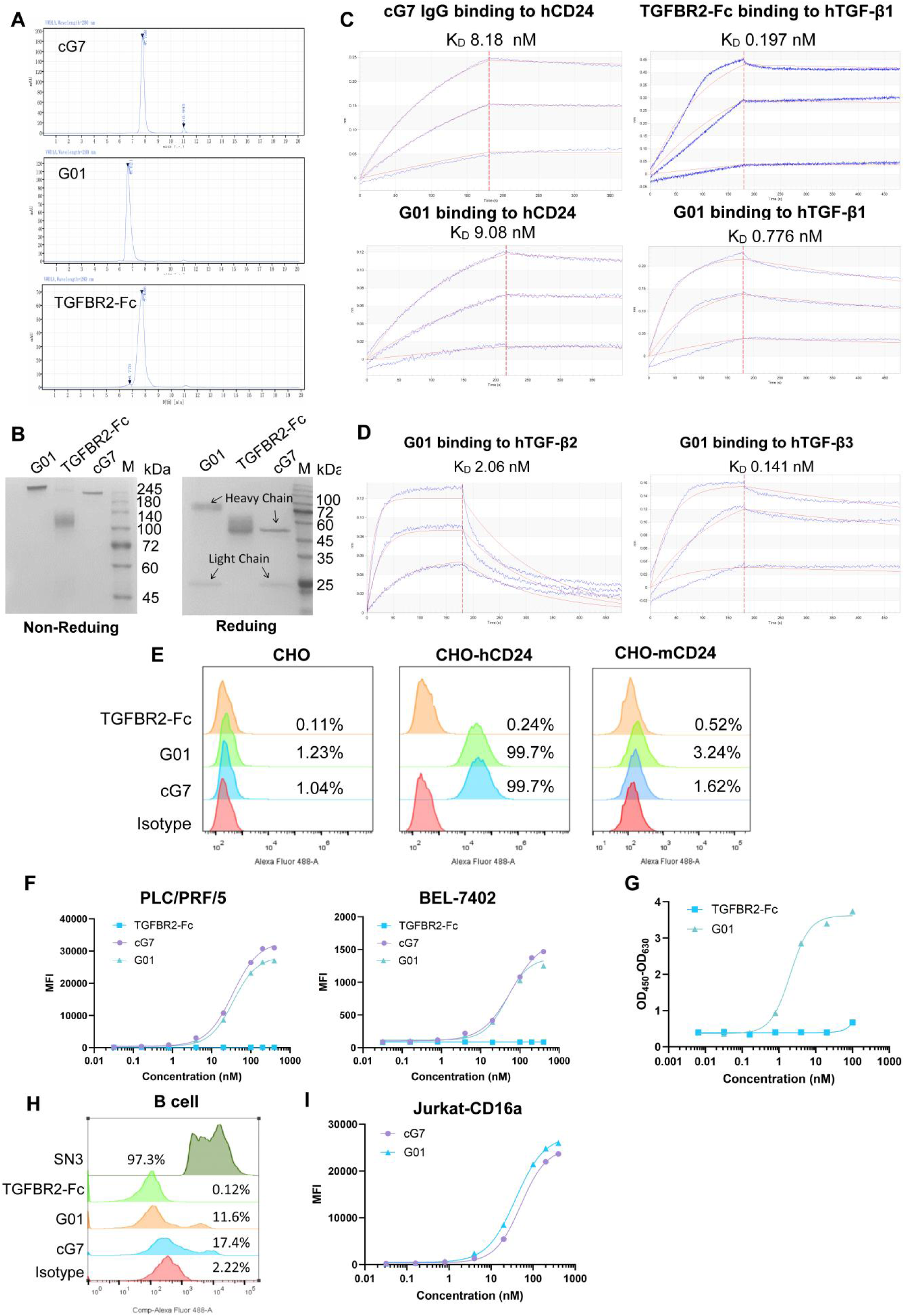
Characterization of G01 in vitro. **(A)** Size-exclusion high-performance liquid chromatography (SEC-HPLC) profiles of G01, cG7, and TGFBR2-Fc. **(B)** SDS-PAGE analysis of G01, cG7, and TGFBR2-Fc under reducing and non-reducing conditions. **(C)** Binding kinetics of cG7, TGFBR2-Fc, and G01 to CD24 and TGF-β1, measured by biolayer interferometry (BLI). **(D)** Binding kinetics of human TGF-β2 and TGF-β3 to G01, measured by BLI. **(E)** Binding of TGFBR2-Fc, cG7 and G01 to human or mouse CD24 overexpressing CHO cells, determined by flow cytometry. **(F)** Binding of G01 to CD24-expressing tumor cells (PLC/PRF/5, BEL-7402), determined by flow cytometry. Curve fitting and EC_50 c_alculation were performed using a sigmoidal 4PL regression model. **(G)** Concurrent binding of G01 to CD24 and TGF-β1 detected by ELISA. Plate-bound hTGF-β1 captured G01, which subsequently binds to mFc-tagged hCD24 antigen, probed by an HRP-conjugated anti-mFc antibody. Curve fitting and EC_50 c_alculation were performed using a sigmoidal 4PL regression model. **(H)** Binding of SN3, TGFBR2-Fc, G01 or cG7 to B cells in human blood. **(I)** Binding of cG7 and G01 to CD16a-expressing Jurkat cells, determined by flow cytometry.

**Fig. S2.**
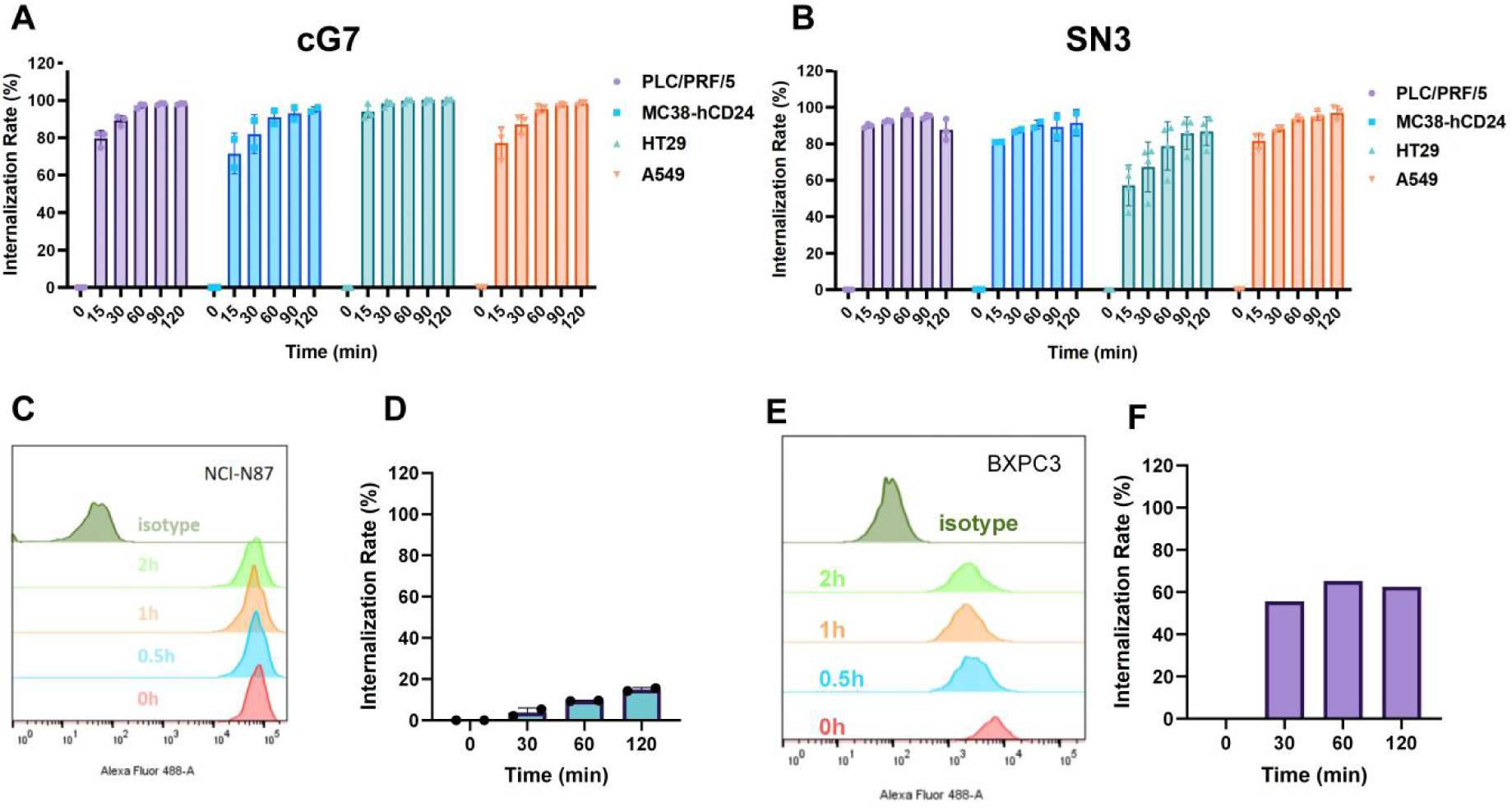
CD24 mediates rapid internalization of associated antibodies. **(A and B)** Kinetics of CD24 monoclonal antibody cG7 (A) and SN3 (B) internalization in PLC/PRF/5, HT29, A549, and MC38-hCD24 cells, quantified by flow cytometry. Data are representative of two independent experiments and presented as mean ± SD. **(C and D)** Kinetics of HER2 monoclonal antibody (Herceptin) internalization in NCI-N87 cells, quantified by flow cytometry. Representative histograms of fluorescence intensity shifts (C) and the time-dependent internalization rate (D) were shown. Data are are representative of two independent experiments and presented as mean ± SD. **(E and F)** Kinetics of Trop2 monoclonal antibody (FDA018) internalization in BxPC3 cells, quantified by flow cytometry. Representative histograms of fluorescence intensity shifts (E) and the time-dependent internalization rate (F) were shown. Internalization rate (%) was calculated based on the mean fluorescence intensity (MFI) using the formula: [MFI (0 h) − MFI (n h)] / [MFI (0 h) − MFI (isotype)] × 100%.

**Fig. S3.**
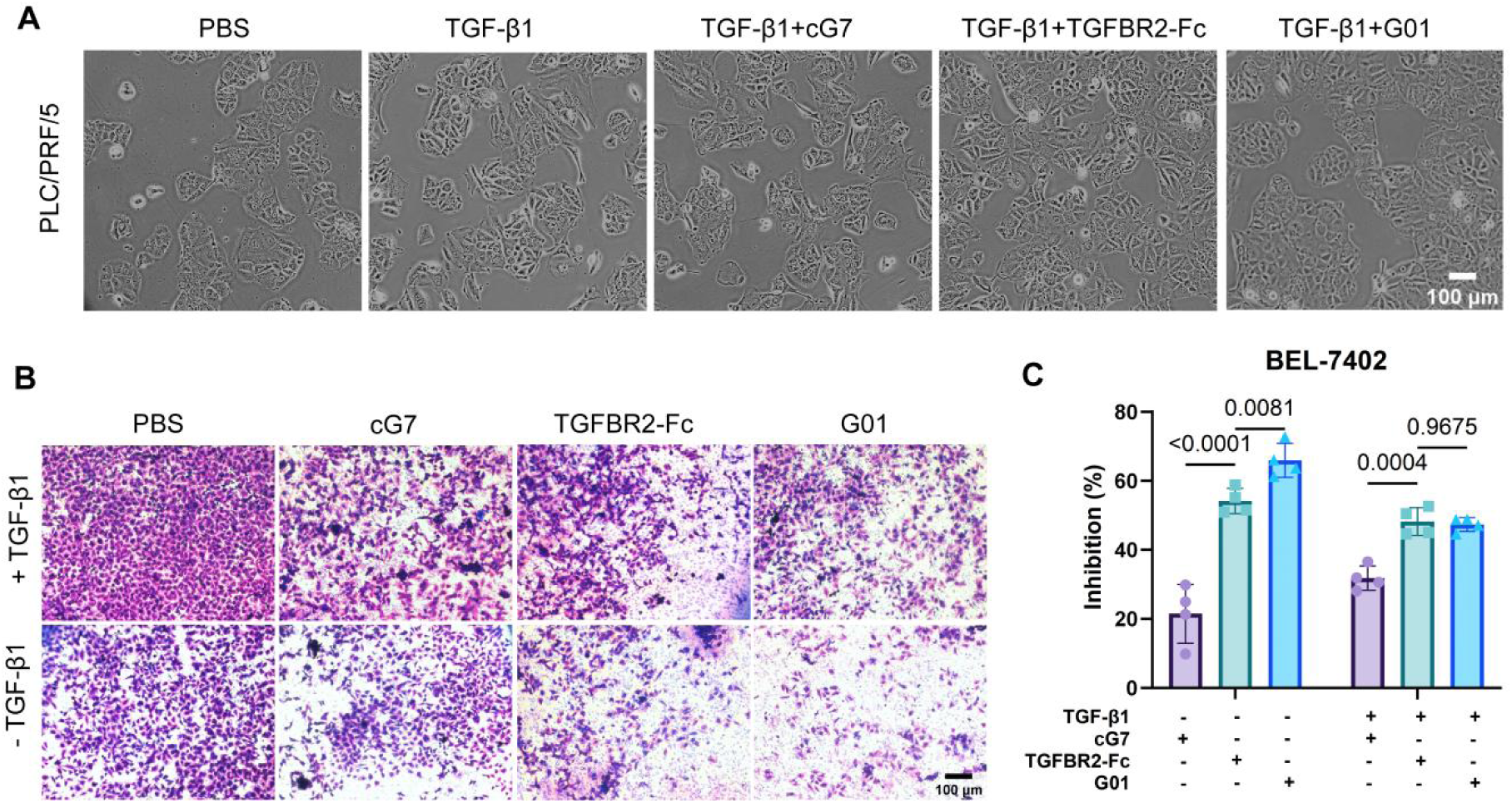
G01 suppresses epithelial-mesenchymal transition (EMT) in tumor cells. **(A)** Representative images showing the morphology of PLC/PRF/5 cells upon different treatments. Scale bar, 100 μm. **(B and C)** Inhibition of BEL-7402 cell invasion by G01. Cells were treated with G01, cG7, or TGFBR2-Fc in the presence of 10 ng/ml human TGF-β1 for 36 hours, stained with 0.1% crystal violet and captured by inverted microscopy (B). The invasion inhibition rate was quantified (C). Scale bar, 100 μm. Data are presented as means ± SEM (n = 3), and statistical significance was determined by one-way ANOVA with multiple comparisons.

**Fig. S4.**
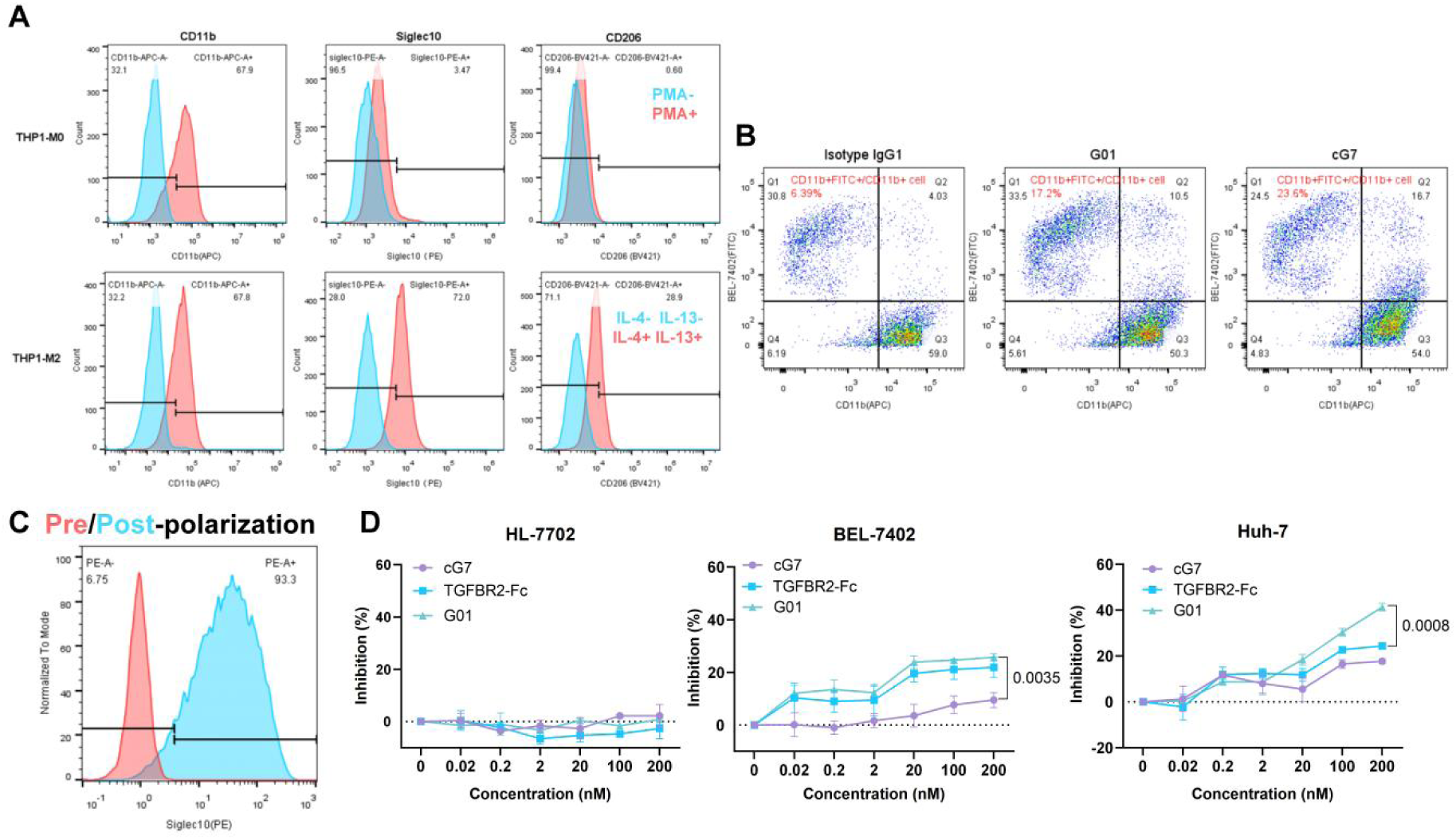
G01 displays Fc-dependent ADCP of CD24^+^ tumor cells and inhibits tumor cell proliferation. **(A)** Representative flow cytometry histograms showing the expression levels of CD11b, Siglec-10, and CD206 on THP-1–derived macrophages. THP-1 cells were first incubated with PMA (100 ng/mL) for 48 hours to generate unpolarized (M0) macrophages, followed by polarization into M2 macrophages using IL-4 (20 ng/mL) and IL-13 (20 ng/mL) in the presence of PMA for an additional 48 hours. **(B)** Representative flow cytometry plots of BEL-7402 phagocytosis by human monocyte-derived macrophages. FITC-labeled BEL-7402 cells were pre-treated with G01, cG7, or an isotype IgG control. Phagocytosis was quantified as the percentage of CD11b^+^ FITC^+^ double-positive cells in CD11b^+^ macrophages. **(C)** Representative flow cytometry histograms evaluating Siglec-10 expression on human monocyte-derived macrophages. Monocytes isolated from PBMCs were differentiated with M-CSF (50 ng/mL) and subsequently stimulated with M2-polarizing cytokines (IL-4 and IL-13, 20 ng/mL each) for 72 hours. **(D)** Impact of G01 on the proliferation of CD24^+^ hepatocellular carcinoma cell lines BEL-7402 and Huh-7 and CD24^-^normal liver cell line HL-7702, assessed by MTT assay. Data are presented as mean ± SD, and statistical significance was determined using two-way ANOVA with Tukey’s post hoc test.

**Fig. S5.**
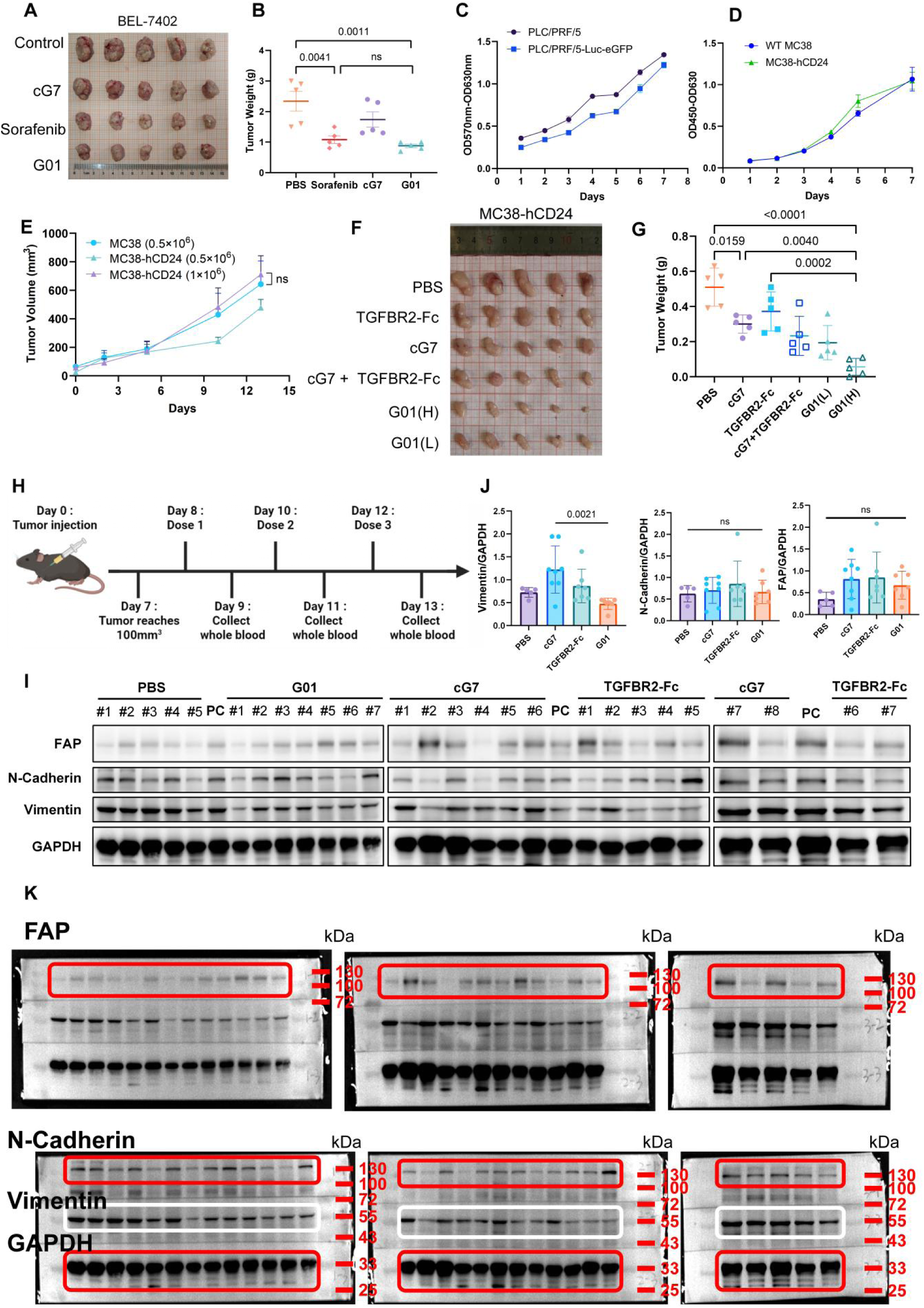
G01 suppresses tumor progression in multiple mouse tumor models. **(A and B)** Antitumor efficacy of G01 in BEL-7402 xenograft model. Representative images of excised tumors (A) and quantification of tumor weights (B) at the study endpoint (day 17). n=5, data are presented as mean ± SD. Statistical significance was determined using one-way ANOVA with Tukey’s post hoc test. **(C)** In vitro proliferation of PLC/PRF/5-Luc cells compared with that of the parental PLC/PRF/5-WT cells, assessed by CCK-8 assay. n=3, data are presented as mean ± SD. **(D and E)** Impact of human CD24 expression on MC38 tumor growth. Proliferation of MC38-hCD24 and MC38-WT cells was evaluated in vitro by CCK-8 assay (D) and in vivo by monitoring tumor growth in syngeneic C57BL/6J mice (E). n=3, data are presented as mean ± SD. Statistical significance was determined using two-way ANOVA with Tukey’s post hoc test. **(F and G)** Antitumor efficacy of G01 in a syngeneic immunocompetent model. Representative images of excised tumors (F) and tumor weights at the study endpoint (day 14) (G) are shown. **(H)** Schematic of the treatment regimen in mice bearing subcutaneous MC38-hCD24 tumors for TGF-β1 detection. Mice bearing established tumors were treated intraperitoneally every 2 days for a total of 3 doses with G01 (n = 8, 20 mg/kg) or equimolar doses of cG7 (n = 8, 16.67 mg/kg). TGFBR2-Fc (n = 8, 8.89 mg/kg) served as a positive control, and PBS (n = 5) was used as a vehicle control. **(I)** Western blot analysis and quantification of fibroblast activation protein (FAP), N-cadherin and vimentin levels in MC38-hCD24 tumors. PC, positive control, was loaded onto every gel as an inter-gel calibrator to correct variability. n = 5 for PBS group, n = 8 for cG7 group, n = 7 for G01 and TGFBR2-Fc groups. **(J)** Quantitative densitometry analysis of the western blots. Protein levels of vimentin was normalized to GAPDH. All densitometry data were quantified using ImageJ software. Data represent mean ± SD, and statistical significance was determined by one-way ANOVA with Tukey’s post hoc test. **(K)** Figure S5I source data. Membranes were cut prior to hybridization with antibodies based on molecular weights. The same membrane was stripped and sequentially reprobed for FAP and N-Cadherin.

**Fig. S6.**
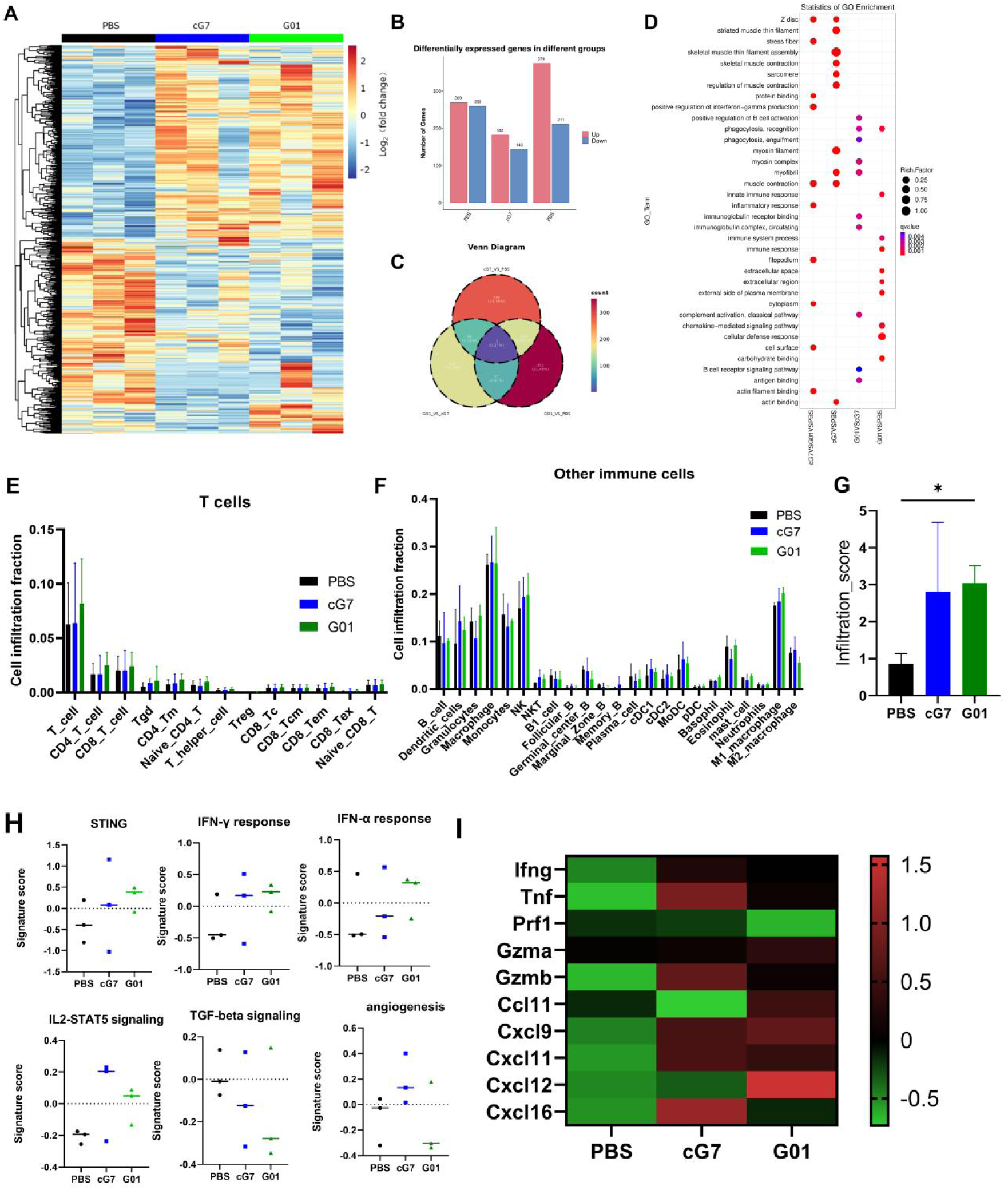
Transcriptomic profiling reveals that G01 treatment remodels the tumor microenvironment to foster innate and adaptive immune activation. RNA-sequencing (RNA-seq) was performed on excised MC38-hCD24 tumors from C57BL/6J mice treated intraperitoneally with PBS, cG7 (8.3 mg/kg), or G01 (10 mg/kg) every 2 days for a total of 5 doses (n = 3). **(A)** Heatmap displaying the expression patterns of all significantly differentially expressed genes (DEGs; defined as P<0.05 and |fold change| > 1.5). Rows represent individual genes, and columns represent individual mice. **(B)** Bar graph quantifying the total number of up and down regulated DEGs in each treatment group. **(C)** Venn diagram illustrating the intersection of DEGs derived from three comparisons: G01 vs. PBS, cG7 vs. PBS, and G01 vs. cG7. **(D)** Bubble plot of Gene Ontology (GO) enrichment analysis. **(E to G)** Evaluation of tumor immune infiltration using ImmuCellAI. Bar charts detailing the estimated infiltration scores for various T cell subtypes (E) and other immune cells (F) (including B cells, NK cells, monocytes, macrophages, neutrophils, and dendritic cells). (G) Overall immune infiltration score. **(H)** Gene Set Variation Analysis (GSVA) signature scores for key immune signaling pathways. **(I)** Heatmap visualizing the transcriptomic profiles of crucial lymphocyte-recruiting chemokines and cytotoxicity-associated effector molecules.

**Fig. S7.**
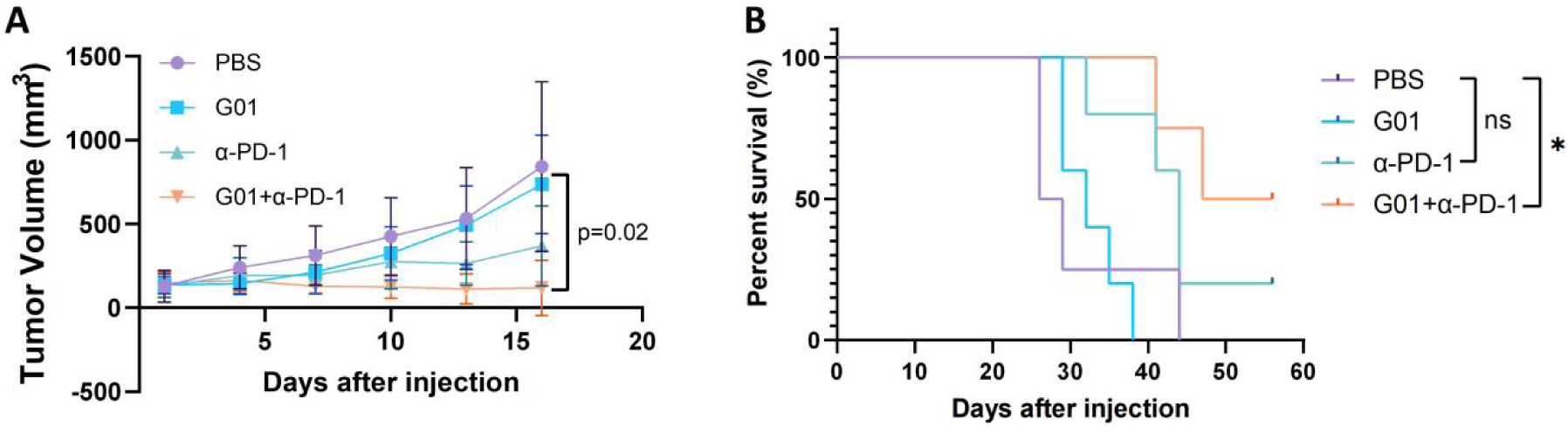
In vivo anti-tumor efficacy of G01 in combination with PD-1 blockade. **(A)** Tumor growth kinetics in the MC38-hCD24 subcutaneous model. Mice bearing established tumors were randomized (n = 5 for G01 and anti-PD-1 groups, n = 4 for PBS and G01+ anti-PD-1 groups) and treated intraperitoneally every three days for a total of 3 doses with G01 (3 mg/kg), anti-PD-1 antibody (RMP1-14, 3 mg/kg) or combination of both when tumor volumes reached 100-150 mm^3^. Tumor volumes were monitored every three days. Data are presented as mean ± SD, and statistical significance was determined by two-way ANOVA. **(B)** Kaplan-Meier survival analysis of tumor-bearing mice. Statistical significance was determined by Log-rank (Mantel-Cox) test.

**Fig. S8.**
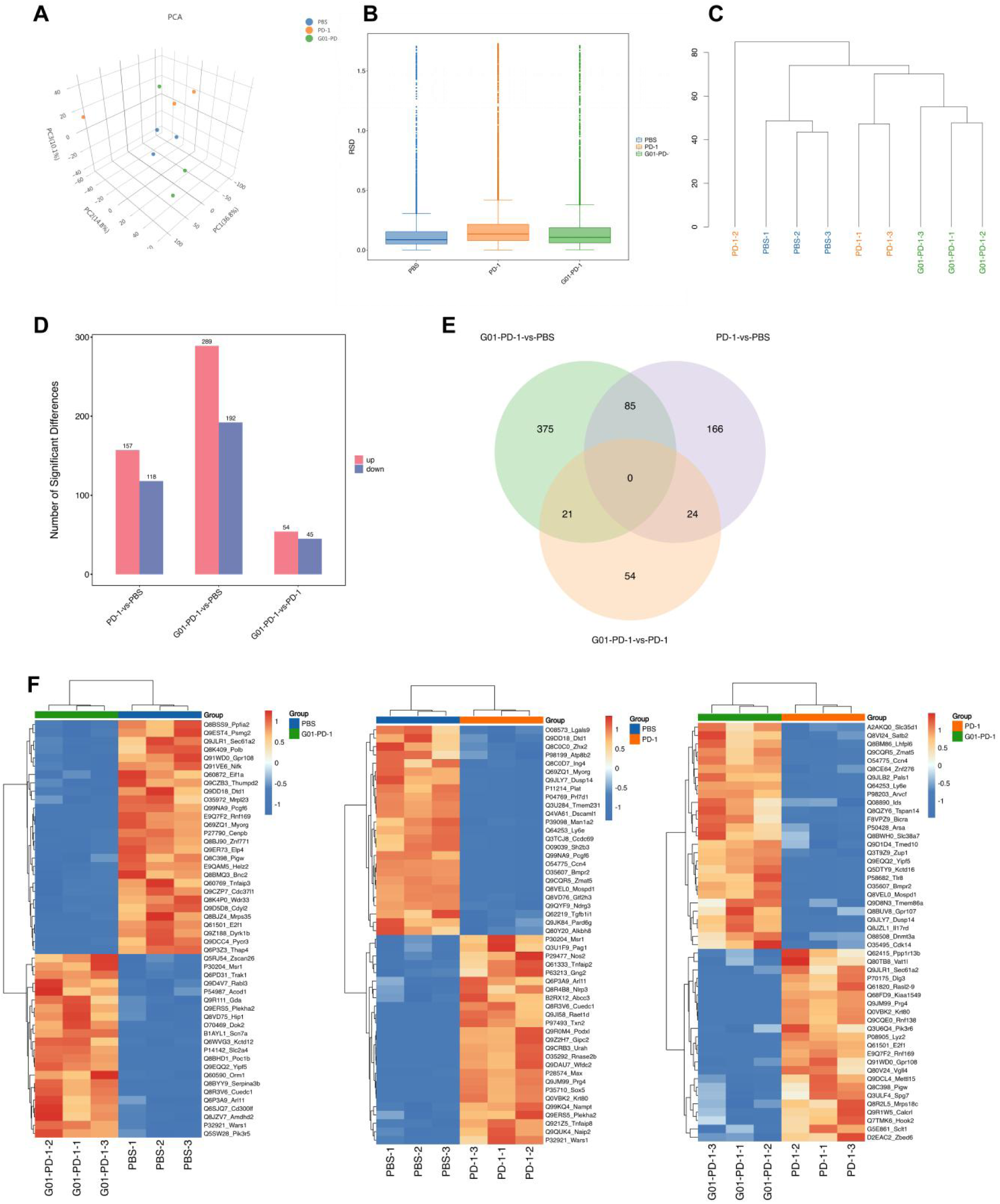
Quality control and quantitative distribution of the DIA-based proteomic profiling. Total proteins were extracted from MC38-hCD24 tumor tissues and analyzed via Data-Independent Acquisition (DIA) mass spectrometry (n = 3 per group). **(A)** Three-dimensional principal component analysis (3D-PCA) illustrating the separation of PBS, anti-PD-1, and G01+anti-PD-1 treatment groups. **(B)** Box plot showing the relative standard deviation (RSD) across samples to evaluate technical and biological reproducibility (RSD = SD/mean). **(C)** Hierarchical clustering analysis (HCA) based on Euclidean distance, depicting the transcriptomic similarity and clustering relationships among individual samples. **(D and E)** Summary of differentially expressed proteins (DEPs; defined as fold change [FC] ≥ 1.2 and P<0.05) across three pairwise comparisons: G01+anti-PD-1 vs. PBS, anti-PD-1 vs. PBS, and G01+anti-PD-1 vs. anti-PD-1. Results are presented as a bar graph indicating the number of upregulated and downregulated proteins (D), and a Venn diagram showing the overlap of DEPs among the indicated groups (E). **(F)** Representative heatmaps displaying the expression patterns of the top 50 DEPs (ranked by fold change) for each pairwise comparison across all biological replicates.

**Fig. S9.**
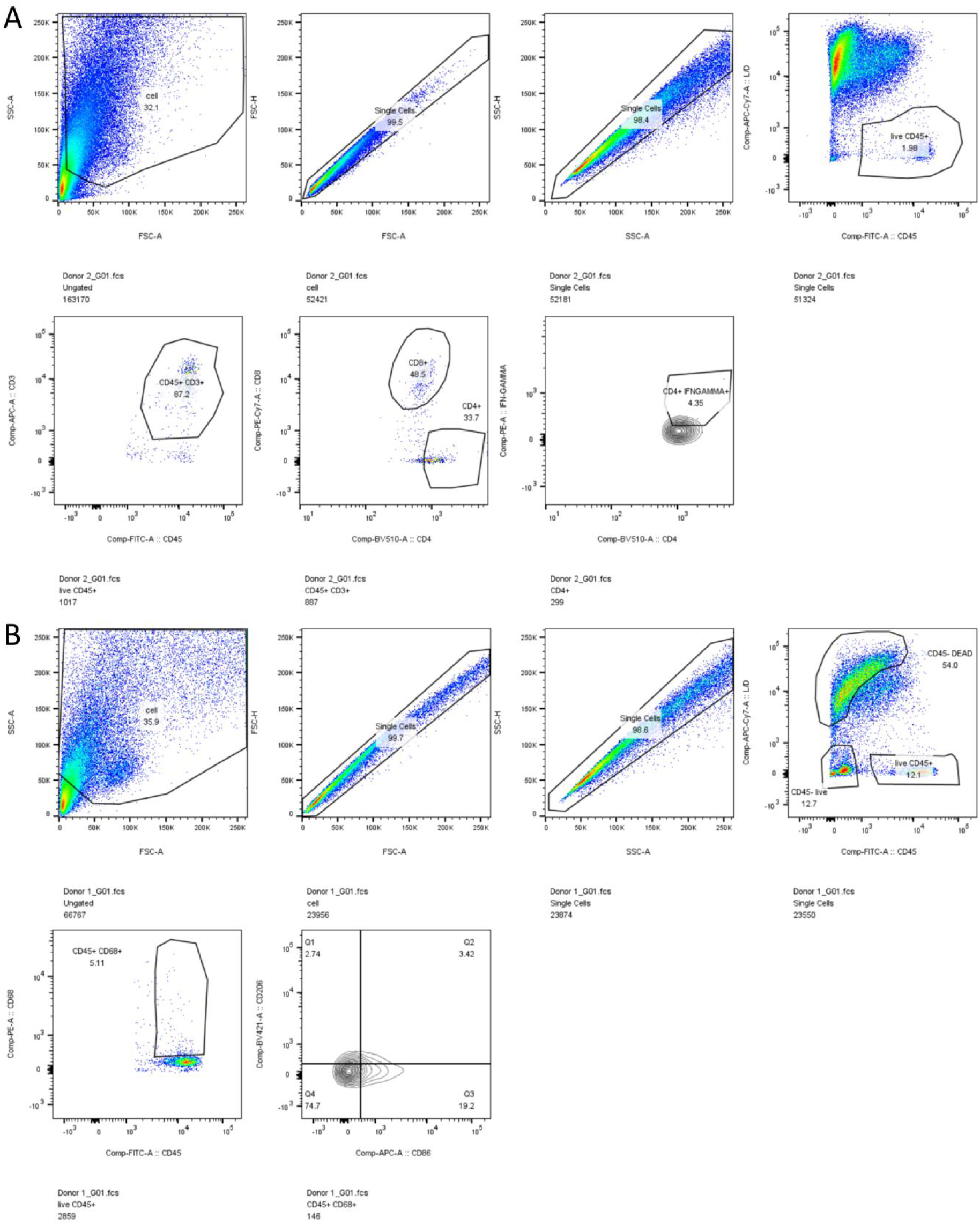
Flow cytometric gating strategies for characterization of tumor-infiltrating immune cells in human PDTFs. Representative flow cytometry plots illustrating the sequential gating hierarchy used to identify T cell (A) and macrophage (B) subsets within PDTFs.

## References and Notes

1. Shen KY, Zhu Y, Xie SZ, Qin LX. Immunosuppressive tumor microenvironment and immunotherapy of hepatocellular carcinoma: current status and prospectives. J Hematol Oncol. 29;17(1):25 (2024).

2. Liu F, Li X, Zhang Y, Ge S, Shi Z, Liu Q, Jiang S. Targeting tumor-associated macrophages to overcome immune checkpoint inhibitor resistance in hepatocellular carcinoma. J Exp Clin Cancer Res. 5;44(1):227 (2025).

3. Wang Z, Wu J, Li Q, Wen X. Overcoming immune resistance in hepatocellular carcinoma: insights into mechanisms, predictive factors, and interventional strategies. Front Immunol. 13;17:1804366 (2026).

4. Kung HC, Zheng KW, Zimmerman JW, Zheng L. The tumour microenvironment in pancreatic cancer - new clinical challenges, but more opportunities. Nat Rev Clin Oncol. 22(12):969–995 (2025).

5. Sherman MH, Beatty GL. Tumor Microenvironment in Pancreatic Cancer Pathogenesis and Therapeutic Resistance. Annu Rev Pathol. 24;18:123–148 (2023).

6. Xiao Z, Todd L, Huang L, Noguera-Ortega E, Lu Z, Huang L, Kopp M, Li Y, Pattada N, Zhong W, Guo W, Scholler J, Liousia M, Assenmacher CA, June CH, Albelda SM, Puré E. Desmoplastic stroma restricts T cell extravasation and mediates immune exclusion and immunosuppression in solid tumors. Nat Commun. 22;14(1):5110 (2023).

7. Massagué J, Sheppard D. TGF-β signaling in health and disease. Cell. 14;186(19):4007–4037 (2023).

8. Thomas DA, Massagué J. TGF-beta directly targets cytotoxic T cell functions during tumor evasion of immune surveillance. Cancer Cell. 8(5):369–80 (2025).

9. Jana S, Jailwala P, Haribhai D, Waukau J, Glisic S, Grossman W, Mishra M, Wen R, Wang D, Williams CB, Ghosh S. The role of NF-kappaB and Smad3 in TGF-beta-mediated Foxp3 expression. Eur J Immunol. 39(9):2571–83 (2009).

10. Papaspyridonos M, Matei I, Huang Y, do Rosario Andre M, Brazier-Mitouart H, Waite JC, Chan AS, Kalter J, Ramos I, Wu Q, Williams C, Wolchok JD, Chapman PB, Peinado H, Anandasabapathy N, Ocean AJ, Kaplan RN, Greenfield JP, Bromberg J, Skokos D, Lyden D. Id1 suppresses anti-tumour immune responses and promotes tumour progression by impairing myeloid cell maturation. Nat Commun. 29;6:6840 (2015).

11. Tauriello DVF, Sancho E, Batlle E. Overcoming TGFβ-mediated immune evasion in cancer. Nat Rev Cancer. 22(1):25–44 (2022).

12. Grauel AL, Nguyen B, Ruddy D, Laszewski T, Schwartz S, Chang J, Chen J, Piquet M, Pelletier M, Yan Z, Kirkpatrick ND, Wu J, deWeck A, Riester M, Hims M, Geyer FC, Wagner J, MacIsaac K, Deeds J, Diwanji R, Jayaraman P, Yu Y, Simmons Q, Weng S, Raza A, Minie B, Dostalek M, Chikkegowda P, Ruda V, Iartchouk O, Chen N, Thierry R, Zhou J, Pruteanu-Malinici I, Fabre C, Engelman JA, Dranoff G, Cremasco V. TGFβ-blockade uncovers stromal plasticity in tumors by revealing the existence of a subset of interferon-licensed fibroblasts. Nat Commun. 9;11(1):6315 (2020).

13. Huang Z, Chen J, Zhu T, Li J, Ng HY, Zhou Y, Gu X, Xu S, Jia R. Cancer-associated fibroblasts in the tumor microenvironment: heterogeneity, crosstalk mechanisms, and therapeutic implications. Mol Cancer. 18;25(1):19 (2025).

14. Giarratana AO, Prendergast CM, Salvatore MM, Capaccione KM. TGF-β signaling: critical nexus of fibrogenesis and cancer. J Transl Med. 26;22(1):594 (2024).

15. Barkal AA, Brewer RE, Markovic M, Kowarsky M, Barkal SA, Zaro BW, Krishnan V, Hatakeyama J, Dorigo O, Barkal LJ, Weissman IL. CD24 signalling through macrophage Siglec-10 is a target for cancer immunotherapy. Nature. 572(7769):392–396 (2019).

16. Bradley CA. CD24 - a novel ‘don’t eat me’ signal. Nat Rev Cancer. 19(10):541 (2019).

17. Dempsey, L.A. Inhibiting phagocytosis. Nat Immunol. 20, 1089 (2019).

18. Nakazono C, Okada S, Tando S, Ishihara S, Shimomura M, Furuya T, Kameyama K, Küffer S, Müller D, Marx A, Teramukai S, Ströbel P, Itoh K, Inoue M. Transforming Growth Factor-β1 as a Biomarker of Malignant Behavior and PD-L1 Expression in Thymic Epithelial Tumors. Cancer Sci. 117(1):181–192 (2026).

19. Ravi, R., Noonan, K., Pham, V., Bedi, R., Zhavoronkov, A., Ozerov, I. V., Makarev, E., Artemov, A. V., Wysocki, P., & Mehra, R. Bifunctional immune checkpoint-targeted antibody-ligand traps that simultaneously disable TGFβ enhance the efficacy of cancer immunotherapy. Nature Communications, 9(1):741–741 (2018).

20. Gulley JL, Schlom J, Barcellos-Hoff MH, Wang XJ, Seoane J, Audhuy F, Lan Y, Dussault I, Moustakas A. Dual inhibition of TGF-β and PD-L1: a novel approach to cancer treatment. Mol Oncol. 16(11):2117–2134 (2022).

21. Lan Y, Yeung TL, Huang H, Wegener AA, Saha S, Toister-Achituv M, Jenkins MH, Chiu LY, Lazorchak A, Tarcic O, Wang H, Qi J, Locke G, Kalimi D, Qin G, Marelli B, Yu H, Gross AW, Derner MG, Soloviev M, Botte M, Sircar A, Ma H, Sood VD, Zhang D, Jiang F, Lo KM. Colocalized targeting of TGF-β and PD-L1 by bintrafusp alfa elicits distinct antitumor responses. J Immunother Cancer. 10(7):e004122 (2022).

22. Li T, Wang X, Niu M, Wang M, Zhou J, Wu K, Yi M. Bispecific antibody targeting TGF-β and PD-L1 for synergistic cancer immunotherapy. Front Immunol. 13;14:1196970 (2023).

23. Lim HY, Heo J, Peguero JA, Ryoo BY, Decaens T, Barlesi F, Moehler MH, Jehl G, Eggleton SP, Bajars M, Gulley JL. Efficacy and safety of bintrafusp alfa in 2 phase I expansion cohorts with advanced HCC. Hepatology. 81(1):32–43 (2025).

24. Spira A, Wertheim MS, Kim EJ, Tan B, Lenz HJ, Nikolinakos P, Rich PL, Jehl G, Machl A, Ito R, Gulley JL, Kopetz S. Bintrafusp Alfa: A Bifunctional Fusion Protein Targeting PD-L1 and TGF-β, in Patients with Pretreated Colorectal Cancer: Results from a Phase I Trial. Oncologist. 28(2):e124–e127 (2023).

25. Yingling JM, Blanchard KL, Sawyer JS. Development of TGF-beta signalling inhibitors for cancer therapy. Nat Rev Drug Discov. 3(12):1011–22 (2024).

26. Teicher BA. TGFβ-Directed Therapeutics: 2020. Pharmacol Ther. 217:107666 (2021).

27. Akhurst RJ. Targeting TGF-β Signaling for Therapeutic Gain. Cold Spring Harb Perspect Biol. 9(10):a022301 (2017).

28. Teixeira AF, Ten Dijke P, Zhu HJ. On-Target Anti-TGF-β Therapies Are Not Succeeding in Clinical Cancer Treatments: What Are Remaining Challenges? Front Cell Dev Biol. 8:605 (2020).

29. Connolly EC, Freimuth J, Akhurst RJ. Complexities of TGF-β targeted cancer therapy. Int J Biol Sci. 8(7):964–78 (2012).

30. Ganesh K, Massagué J. TGF-β Inhibition and Immunotherapy: Checkmate. Immunity. 48(4):626–628 (2018).

31. Fadok VA, Bratton DL, Konowal A, Freed PW, Westcott JY, Henson PM. Macrophages that have ingested apoptotic cells in vitro inhibit proinflammatory cytokine production through autocrine/paracrine mechanisms involving TGF-beta, PGE2, and PAF. J Clin Invest. 101(4):890–8 (1998).

32. Werfel TA, Cook RS. Efferocytosis in the tumor microenvironment. Semin Immunopathol. 40(6):545–554 (2018).

33. Noy R, Pollard JW. Tumor-associated macrophages: from mechanisms to therapy. Immunity. 41(1):49–61 (2014).

34. Seo W, Jerin C, Nishikawa H. Transcriptional regulatory network for the establishment of CD8^+^ T cell exhaustion. Exp Mol Med. 53(2):202–209 (2021).

35. Wherry EJ, Kurachi M. Molecular and cellular insights into T cell exhaustion. Nat Rev Immunol. 15(8):486–99 (2015).

36. Golestanifar A, Seyedtaghia MR, Mousavi P. Integrated bulk and single-cell transcriptomic profiling reveals NECTIN-TIGIT interaction underlies T cell exhaustion in papillary thyroid carcinoma. Clin Exp Med. 26(1):13 (2025).

